# SPT5 regulates Pol II pausing and elongation in different ways at early versus late embryonic stages

**DOI:** 10.1101/2025.09.06.674612

**Authors:** Alessandro Dulja, Marvin Mayer, Niklas Engel, Arkadiy K. Golov, Mattia Forneris, Yacine Kherdjemil, Songjie Feng, Rebecca R. Viales, Eileen E.M. Furlong

## Abstract

Transcription involves a cycle of initiation, pausing, elongation, and termination. SPT5 regulates both promoter proximal pausing and elongation, but how it orchestrates both steps during dynamic changes in gene expression remains unclear. Here, using *Drosophila* embryogenesis, we show that pausing both precedes and follows gene expression, while active transcription is accompanied by pause release. Optogenetic rapid depletion of SPT5 from the nucleus uncovered different sensitivities at different developmental stages. In early embryogenesis, SPT5 depletion caused a downstream shift in Pol II pausing to the +1 nucleosome, resulting in defective elongation and early termination. In late embryogenesis, it led to both up- and downregulation of expression, depending on the genes’ transcriptional and pausing state – upregulation is caused by pause release while downregulation is due to defective elongation. These changes are intensified when genes are increasing or decreasing their transcriptional state, indicating that SPT5 contributes to fine-tuning dynamic changes in gene expression.

## INTRODUCTION

Transcription comprises a cycle of precisely regulated steps, each contributing to the overall transcriptional output. Initiation involves the recruitment of general transcription factors and RNA polymerase II (Pol II) to the promoter, forming the pre-initiation complex (PIC)^1^. At many genes, after transcribing a short stretch of 30-80 nucleotides, the complex undergoes promoter-proximal pausing that results in either early termination or the progression into productive elongation^2,3^. Rather than involving stable stalling as initially assumed, recent studies indicate that pausing is a dynamic process characterised by high rates of Pol II turnover^4–6^. Pausing is therefore a crucial regulatory step representing a kinetic bottleneck in the transcription cycle. Given that, pause release, in addition to Pol II recruitment and initiation, are key steps that transcription factors regulate to activate gene expression^3^.

Pausing is mainly induced by negative elongation factor (NELF) and the DRB-sensitivity-inducing factor (DSIF), the latter being composed of two subunits; suppressor of Ty5 (SPT5) and suppressor of Ty4 (SPT4)^7–9^. NELF binds to and stabilises the pre-formed DSIF/Pol II complex^8,10^ In this configuration, NELF and DSIF are unphosphorylated, while Pol II is phosphorylated at the Ser5 residues of its C-terminal domain (CTD) repeats. The pausing phase resolves into productive elongation when multiple components of the paused NELF/DSIF/Pol II complex are phosphorylated by the CDK9 subunit of the positive transcription elongation factor b (P-TEFb) complex^11^. Phosphorylation of Pol II at the Ser2 residues of its CTD repeats makes it highly processive and thus elongation competent^12^. Phosphorylated NELF dissociates from the complex, where it is replaced by the elongation factors PAF and SPT6^13,14^, while phosphorylation of SPT5 converts it into a positive elongation factor that remains associated with elongating Pol II^8,15^. SPT5 therefore acts as both a pausing factor at the promoter and an elongation factor along the gene body, depending on its phosphorylation status. The function of SPT5 in elongation is conserved across all domains of life and it is considered a universal elongation factor^16,17^, while its role in pausing appears to have evolved more recently, coinciding with the evolution of increased regulatory complexity^18^. The smaller DSIF subunit SPT4 plays a more minor role^19^.

Much of this knowledge has come from biochemical and structural studies, including *in vitro* reconstitution experiments to assess Pol II activity in the presence of the different pausing factors^3,18^. In contrast, functional studies using classic loss-of-function mutants have been hindered due to the essential role of these factors. However, recent acute protein degradation methods^20,21^ can overcome these issues, and their application to transcriptional regulators led to surprising results. In the human DLD-1 cancer cell line, for example, the rapid depletion of NELF did not result in higher productive elongation as expected^22^. Instead, Pol II moved to a second pausing site about 50 bp more downstream of the canonical one, corresponding to the +1 nucleosome^22^. Directly assessing the impact of SPT5 depletion on transcription is more difficult, as it also plays a role in Pol II stabilisation. In the same cell line, the acute degradation of SPT5 caused the degradation of the main Pol II subunit RPB1 and thus led to a global decrease in Pol II chromatin occupancy^23^, complicating the interpretation of the results. The proteasomal degradation of RPB1 is mediated through VCP/p97^24^, an enzyme involved in the segregation of ubiquitylated proteins from their interaction partners. To overcome this, a recent study combined SPT5 depletion with pharmacological VCP inhibition to stabilise Pol II^24^, which resulted in Pol II pausing at the +1 nucleosome, phenocopying the depletion of NELF. This supports previous reports that the +1 nucleosome is a strong barrier for Pol II elongation^25^ and represents a second, more downstream pausing site^26^. In keeping with this, a second study in human cell lines used a slower, partial depletion of SPT5 (that kept RPB1 intact for the first hours), which also detected a shift in Pol II occupancy to a region more downstream of the TSS^27^.

In most cell culture models – unless the cells are induced into differentiation^28^ – transcription is at steady state. It remains unclear what role SPT5 has during dynamic gene expression changes when genes are switching on and off, and how these changes are modulated by either Pol II pausing or elongation. To address these questions, we used embryogenesis as a model system, which is characterised by highly dynamic transcriptional changes, especially in *Drosophila* which completes embryogenesis in less than 24 hours. Perturbing pausing in living embryos is very challenging, as loss-of-function mutants removing components of the transcriptional machinery often led to a block in oogenesis or embryogenesis at very early stages. The only genetic evidence for the requirement of SPT5 in *Drosophila* embryos is from a missense mutation in the C-terminal domain (W049), which is homozygous lethal and likely reduces SPT5 activity, leading to the misexpression of some genes in the early blastoderm embryo^31^. A 48-h RNAi knockdown of SPT5 in a *Drosophila* cell line caused a genome-wide reduction in Pol II occupancy and transcription^32^. However, in light of the recent mammalian studies^24,33^, it is unclear if this is mediated through the destabilisation of Pol II. Auxin, or other targeted degradation systems have not been applied in *Drosophila* due to the embryos’ protective impermeable vitelline membrane. Here, we solved these issues using a recently developed optogenetic method, iLEXY^34^, to induce extremely rapid (within minutes) and reversible nuclear depletion of SPT5 at specific developmental time windows after blue light exposure.

In *Drosophila*, Pol II pausing has mostly been described at early blastoderm stages of embryogenesis, when it appears to be widespread^29,30^. To assess pausing throughout all stages of embryogenesis, we first generated a timecourse of SPT5 occupancy and measured Pol II pausing across diverse stages. SPT5 occupancy changes throughout embryogenesis reflecting dynamic changes in gene expression. Pausing is established before gene expression, while active transcription is accompanied by pause release, followed by the re-establishment of Pol II pausing when the gene is switching off. To assess the functional role of SPT5, we applied the iLEXY system^34^ at different stages inducing rapid, but not complete (∼50%) depletion of SPT5, which avoids the degradation of Pol II. Our results uncovered different molecular phenotypes after SPT5 depletion at different stages, which surprisingly uncovered a much higher sensitivity for embryonic viability at later stages. In early embryos, SPT5 depletion caused a downstream shift of Pol II pausing to the +1 nucleosome, accompanied by defective elongation and early transcriptional termination. Interestingly, this switches in late embryogenesis, when SPT5 depletion caused Pol II release into the gene body, without any stalling at the +1 nucleosome. This resulted in either upregulation of gene expression through pause release into productive elongation or downregulation through impaired elongation, depending on a gene’s pausing index and transcriptional state.

## RESULTS

### SPT5 occupancy changes during embryogenesis reflecting dynamic gene expression

To characterise the role of SPT5 during embryogenesis, we used CRISPR-Cas9-mediated homology-directed repair to tag the endogenous *spt5* gene at the C-terminus with a HA-iLEXYs tag. As there are no commercial antibodies directed against *Drosophila* SPT5, the HA tag allows us to examine the occupancy of SPT5 during embryogenesis for the first time, while the iLEXYs module should enable the nuclear depletion of SPT5 from in response to blue light^34^. The *spt5-HA-iLEXYs* line is homozygous viable and fertile under safelight conditions (“dark”), indicating that the tag does not significantly impact the protein’s function.

To determine SPT5 occupancy, we performed CUT&Tag (using an anti-HA antibody) on staged embryo collections from the *spt5-HA-iLEXYs* line kept in the dark. We chose five 2-hour time windows spanning almost all stages of embryogenesis (**Fig. 1A**). The biological replicates are highly correlated, attesting to the quality of the data (**Fig. 1B**). The correlation analysis also revealed an interesting shift at 10-12 h after egg laying (a.e.l.), with the earliest two time points (2-4 and 6-8 h) and latest two time points (14-16 and 18-20 h) being more correlated to each other, suggesting that 10-12 h (the beginning of terminal differentiation) is a transition point in SPT5 function (**Fig. 1B**).

To explore the relationship between the temporal changes in SPT5 occupancy and gene expression, we used a matched time-course of RNA-seq^35^ and applied k-means (k = 3) clustering to group genes based on both their dynamic changes in SPT5 binding (total signal from 250 bp upstream of the TSS to the TES) and gene expression throughout embryogenesis (**Fig. 1C, Table S1**), which captures the three main expression trends of interest – increasing, decreasing, or transient expression across embryonic development. Cluster 1 and cluster 3 represent early or late genes that decrease or increase in expression during embryogenesis, respectively, and in both cases SPT5 binding and RNA levels show coordinated temporal changes, either decreasing or increasing together (**Fig. 1C**). Cluster 2 genes are more transiently expressed in mid embryogenesis, peaking at 10-12 h or 14-16 h, and are often pre-bound by SPT5 in the time-point before they are expressed (**Fig. 1C**). This revealed that the dynamic changes in SPT5 occupancy are highly correlated with the dynamic changes in gene expression, which although expected had not been observed during embryogenesis before.

As SPT5 functions as both a pausing and an elongation factor, for each cluster we investigated the distribution of SPT5 binding at the promoter and along the gene body over the five analysed time windows (**Fig. 1D**). Genes that decrease in their expression (cluster 1) show a drastic reduction of SPT5 signal in their gene bodies (indicating a reduction in productive elongation) but retain a strong promoter signal even at the last two time points when the genes decrease in expression (**Fig. 1D, E**). Genes that increase in expression (cluster 3), already have SPT5 at their promoter before the beginning of transcription, which gradually increases together with the gene body signal until 18-20 h, in keeping with their increasing RNA signal (**Fig. 1D, E**). Cluster 2 genes (transiently expressed) have combined features of the other two clusters, generally showing SPT5 promoter binding both before and after their expression, while the gene body signal only transiently increases concomitantly with their transient transcription. For example, the *kni* (cluster 1) and *col4a1* (cluster 3) genes are more highly expressed in early or late stages of embryogenesis, respectively (**Fig. 1E**). While the SPT5 gene body signal correlates with expression level, the SPT5 promoter signal is maintained for many hours after *kni* is no longer expressed (and is actually unchanged even at the end of embryogenesis (18-20 h)) and appears hours before *col41a* is expressed (**Fig. 1E**). Similarly, for the *vvl* gene (cluster 2), SPT5 is already present at the promoter at 2-4 h when the gene is not expressed, and then increases along the gene body concomitantly with transcription and is later retained at the promoter as transcription decreases.

SPT5 occupancy therefore changes during embryogenesis reflecting dynamic gene expression. While the SPT5 gene body signal matches ongoing transcription, its promoter occupancy appears somewhat uncoupled. This likely reflects SPT5’s different roles in promoter-proximal pausing and transcriptional elongation, which we explore below.

**Figure 1:**
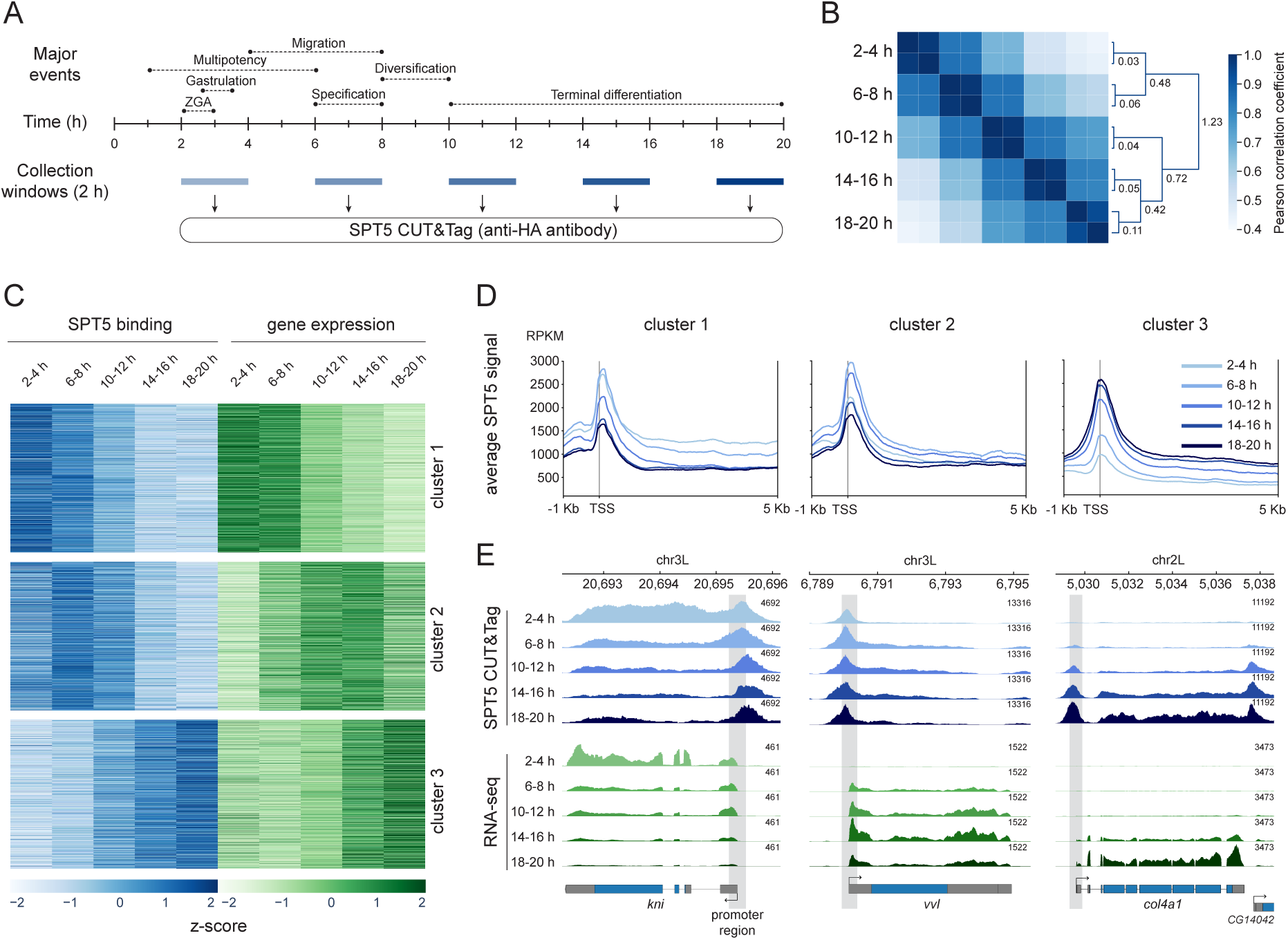
SPT5 occupancy changes during embryogenesis reflecting dynamic gene expression. **(A)** A simplified timeline of *Drosophila* embryogenesis, highlighting major developmental events. The collection windows (blue) for SPT5 CUT&Tag are indicated. **(B)** Pearson correlation heatmap of SPT5 binding across replicates at different developmental time windows considering all *Drosophila* genes. SPT5 binding for each gene was calculated from 250 bp upstream of the TSS to the TES. The dendrogram shows hierarchical clustering, with the sample order reflecting their similarity in binding and branch lengths the relative distance between clusters. **(C)** Heatmap of k-means clusters, showing the dynamics of SPT5 binding (blue) and RNA expression (green). Clustering was performed on genes that are expressed (FPKM > 1) and bound by SPT5 (total signal > 500 RPKM between 250 bp upstream of the TSS to the TES) at one or more time point. Genes in both datasets were normalised using z-scores. Cluster 1: N = 3578; cluster 2: N = 1960; cluster 3: N = 2502. **(D)** Metaplots showing average profiles of SPT5 binding for each gene cluster from (C) over five time windows. **(E)** Genome tracks of SPT5 binding (blue) and RNA expression (green) at five embryonic time windows for *kni* (cluster 1), *vvl* (cluster 2), and *col4a1* (cluster 3). Shaded grey areas indicate the promoter region (TSS ± 250 bp).

### Pausing both precedes and follows gene expression, while active transcription is associated with pause release

As Pol II pausing has been mostly studied during early stages of *Drosophila* embryogensis^29,36–39^, we used our time-course of SPT5 occupancy to investigate how pausing changes across embryogenesis, and how that relates to dynamic gene expression.

Since SPT5 acts both as a promoter pausing and elongation factor, we reasoned that the ratio between its promoter and gene body signal should provide a measure of pausing, similar to a pausing index^40^ (**Fig. 2A**). Before calculating the ratio, the promoter and gene body SPT5 signal were normalised for gene length (see Methods). For each of the five analysed time points, we calculated the SPT5 PI for every gene bound by SPT5 (defined as a total RPKM SPT5 signal of > 500 (from 250 bp upstream of the TSS to the TES)), removing genes shorter than 800 bp to ensure a robust ratio. For each time point, genes were categorised into three classes: (i) bottom 25% (first quartile) – SPT5 PI generally < 1 (**Fig. 2B**), indicating little SPT5 accumulation at their promoter (i.e. not paused), as visible in their SPT5 CUT&Tag metaplots (**Fig. S1A**), (ii) top 25% (fourth quartile) – SPT5 PI generally > 3 (**Fig. 2B**), with strong SPT5 promoter signal (**Fig. S1A**), and (iii) middle 50% (corresponding to the second and third quartile, between 25-75%) (**Fig. 2B, Table S2**).

To validate our SPT5 PI as an indicator of pausing, we mapped the 3’ end of nascent RNA molecules at single-nucleotide resolution using precision nuclear run-on (PRO-seq)^29,41^ at 3-4 and 18-20 h, overlapping the earliest and latest time window analysed, respectively. At both time points, the average PRO-seq signal of the bottom 25% SPT5 PI genes is equally distributed between the promoter and the gene body (**Fig. 2C**), in contrast to the middle 50% and top 25% SPT5 PI genes, which have an increasingly stronger promoter signal that peaks at ∼60 bp downstream of the TSS (**Fig. 2C**), compatibly with the pausing site. Quantitatively, the pausing index calculated from PRO-seq data increases from a median of ∼0.6 for the bottom 25% genes to 2.8 and 3.4 for the top 25% SPT5 PI genes at 2-4 and 18-20 h, respectively (**Fig. S1B**), indicating a high concordance between PI calculated by PRO-seq and SPT5 CUT&Tag.

Next, we investigated how pausing relates to gene expression. The PRO-seq signal varies dramatically among the three SPT5 PI classes only around the pausing site, while it is comparable within the gene body (**Fig. 2C**). In keeping with this, the transcriptional output as measured by RNA-seq is comparable among the three pausing classes (**Fig. 2D**), and the level of Pol II promoter pausing does not correlate with total RNA abundance (**Fig. 2E**).

To explore how pausing relates to dynamic changes in gene expression over development, we looked at changes in SPT5 PI for genes that decrease or increase in expression throughout embryogenesis (clusters 1 and 3 from Fig. 1C, respectively). For both clusters, the SPT5 PI shows a clear bimodal distribution at 18-20 h (**Fig. S2A**). We therefore split the distribution at this latest time point into two populations - roughly those that are continuously increasing or transiently increasing (going up and then down) in their PI over time - and followed how their PI changed from early (2-4 h) to late (18-20 h) stages (**Fig. S2B**). Interestingly, this revealed that genes that decrease (cluster 1) or increase (cluster 3) in expression can both be accompanied by either a steady increase in PI over time or by a transient increase that peaks at mid embryogenesis (**Fig. 2F, S2B**). For genes that decrease in expression (cluster 1), the steady increase in pausing is explained by the decrease in their SPT5 gene body signal over time, while the promoter peak is retained (**Fig. 2F**, left). For the second population (decreased PI by the end of embryogenesis), a similar mechanism can explain the transient increase in pausing until 10-12 h, while the subsequent decrease is likely due to the disassembly of the transcriptional machinery, as indicated by their reduced SPT5 promoter binding (**Fig. 2G** top right, compared to top left). For genes that increase in expression (cluster 3), the continuous increase in pausing is due to the increase in SPT5 promoter binding over time relative to their gene body signal (**Fig. 2G**, bottom left). This likely indicates that the rate of initiation (i.e., the recruitment of the transcriptional machinery) increases over time more than the rate of pause release, explaining their concomitant increase in both pausing and expression. This is compatible with previous data showing that pausing inhibits new initiation^42^. For the second population (with reduced PI at 18-20 h), pausing increases in early embryogenesis suggesting that the machinery is recruited in a paused state, and then pausing starts to decrease concomitantly with the increase in transcription (**Fig. 2G**, bottom right). The increase in transcription for these genes is therefore likely due to pause release, as indicated by their steeper increase in gene body compared to promoter SPT5 signal (**Fig. 2G** bottom right, compared to bottom left). These data indicate that pausing is generally established before the beginning of transcription, while active transcription is accompanied by pause release into productive elongation, which for some genes is accompanied by higher initiation at the promoter, and then pausing is resumed as transcription ends.

Finally, to investigate how the promoters of genes associated with different levels of pausing are regulated, we performed CUT&Tag at 3-4, 10-12, and 18-20 h for the pausing factor NELF as well as serine 5 and serine 2 phosphorylated Pol II, (“pSer5” and “pSer2” hereafter) to capture promoter-proximal (initiating/paused) and elongating forms of Pol II, respectively. The average NELF promoter signal is almost negligible for the bottom 25% PI genes and steadily increases for genes that belong to the middle 50% and top 25% PI classes (**Fig. 2H**). Quantitatively, the percentage of genes whose promoter is significantly bound by NELF increases from less than 20% to ∼40% and ∼80% across the three SPT5 PI classes (**Fig. S2D**). Similarly, Pol II (both pSer5 and pSer2) shows an increasingly higher promoter accumulation that scales with the levels of pausing (**Fig. 2H**). This high concordance between NELF and SPT5/Pol II occupancy is consistent with NELF’s role to stabilise pausing by binding to the SPT5/Pol II complex^8,10^. Of note, the majority of genes are either always bound or always not bound by NELF across the three analysed time windows (**Fig. 2I**), suggesting that the level of SPT5/Pol II, and thus NELF, binding is promoter specific and likely genetically encoded. In agreement, motifs previously associated with pausing like the pause button (PB), downstream promoter element (DPE) and initiator (Inr) motifs^3,43^, are enriched among the genes’ promoters that are constitutively highly paused (SPT5 PI > 3) throughout embryogenesis. Instead, the TATA element is more enriched among the constitutively lowly paused genes (SPT5 PI < 1) (**Fig. S2E**). Intriguingly, genes that are constitutively highly paused (SPT5 PI > 3) throughout embryogenesis are longer than modestly and weakly paused genes, with a median gene length of 9.1 kb compared to 2.5 and 1.9 kb, respectively (**Fig. S2F**). This suggests a selection of highly paused promoters for longer genes, again hinting at a genetic component of pausing.

Taken together, using SPT5 occupancy to measure pausing over embryogenesis revealed that i) the level of pausing (a gene’s pausing index) is uncoupled from its level of expression and that ii) pausing both precedes and follows the expression of a gene, while active transcription is accompanied by pause release and in some cases increased initiation. Lowly paused genes have little SPT5/Pol II accumulation and thus no NELF binding at their promoter, while the opposite is true for highly paused genes. To test the causal role of SPT5 in these processes, we next perturbed SPT5 function.

**Figure 2:**
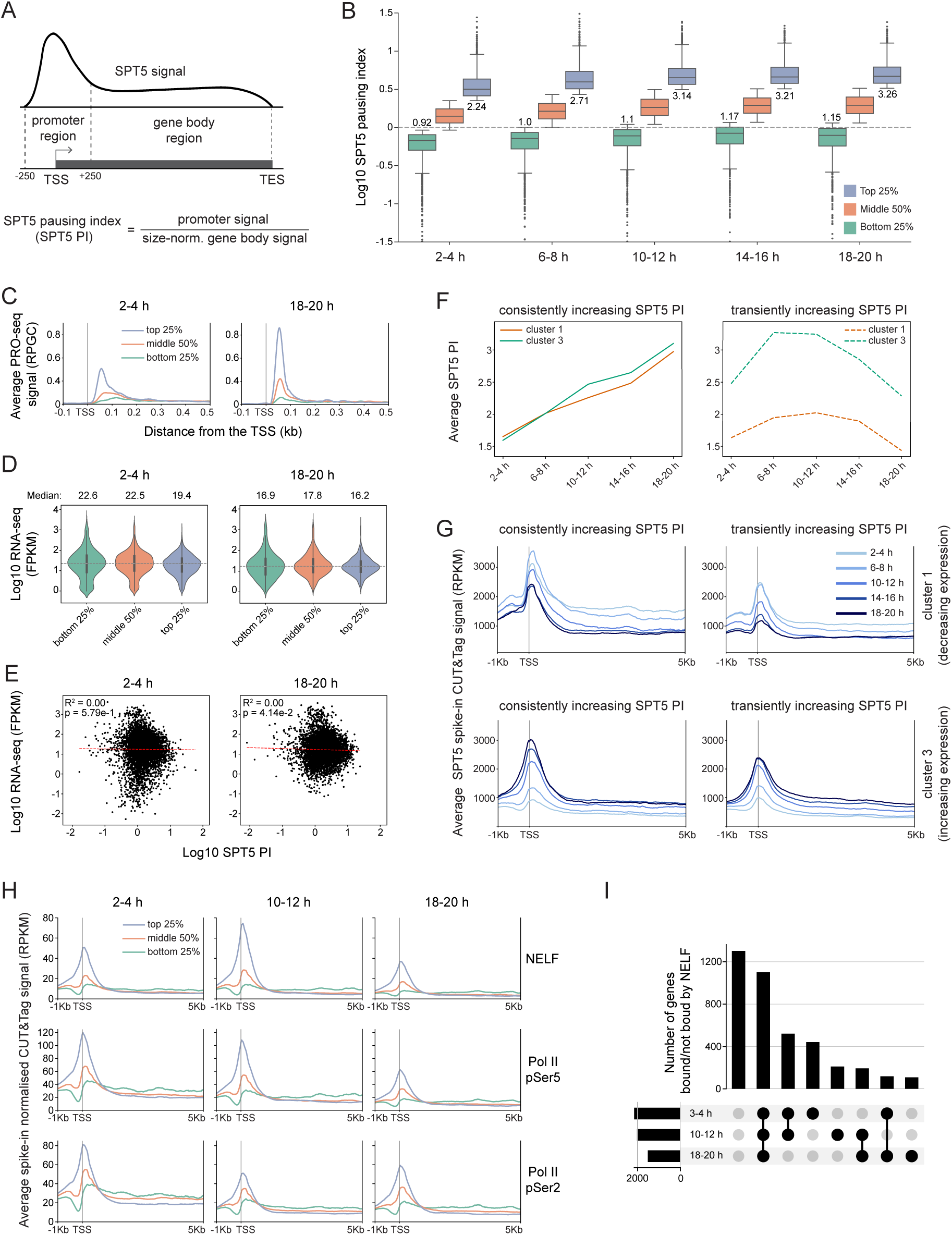
Pausing both precedes and follows gene expression, while active transcription is associated with pause release. **(A)** Schematic of the SPT5 pausing index (PI), calculated as the ratio of promoter signal (TSS ± 250 bp) over the size-normalised gene body (from +250 bp to the TES) signal (see Methods). **(B)** Boxplot showing the SPT5 PI distribution of the three classes at the five analysed time windows. Genes with a total (promoter plus gene body) SPT5 signal < 500 RPKM were deemed unbound and excluded. 2-4 h: N = 5567; 6-8 h: N = 5671; 10-12 h: N = 5806; 14-16 h: N = 5527; 18-20 h: N = 5555. **(C)** Average PRO-seq signal for the three SPT5 PI classes. SPT5 PI classes from 2-4 h embryos were matched with PRO-seq from 3-4 h embryos. Top 25%: N = 1392; Middle 50%: N = 2783; Bottom 25%: N = 1392. For 18-20 h, both datasets were generated from 18-20 h old embryos. Top 25%: N = 1389; Middle 50%: N = 2777; Bottom 25%: N = 1389. **(D)** Violin plots of time-matched expression levels from RNA-seq^50^ or the three SPT5 PI classes at 2-4 h and 18-20 h of embryogenesis. Median values (non-log) shown above. **(E)** Scatterplots showing the correlation between expression (RNA-seq) and SPT5 PI at 2-4 h and 18-20 h of embryogenesis. 2-4 h: N = 5567; 18-20 h: N = 5555. **(F)** Average SPT5 promoter index at the five analysed time windows for genes consistently increasing or transiently increasing in their PI in cluster 1 and 3 from Fig. S2B (cluster 1, consistently increasing: N = 1349; transiently increasing: N = 2229. Cluster 3, steadily increasing: N = 761; transiently increasing: N = 1741). **(G)** Metaprofiles of SPT5 signal for genes consistently increasing or transiently increasing in their PI in cluster 1 and 3 (gene numbers as in (F)). **(H)** Average, normalised spike-in CUT&Tag signal for NELF, Pol II pSer5, and Pol II pSer2 for the three indicated time windows. For SPT5 PI classes, the following CUT&Tag data was used: for 2-4 h, data from 3-4 h embryos (corresponding to Fig. 4 dark control), for 10-12 and 18-20 h time matched data from Fig 5 dark control was used. **(I)** UpSet plot showing gene sets categorised by the presence (black) or absence (grey) of NELF binding at their promoters across the three time windows. Only genes with detected SPT5 signal (> 500 RPKM between 250bp upstream of the TSS to the TES) at all 3 timepoints were included (N = 3959).

### Optogenetic nuclear depletion of SPT5 decreases its binding to chromatin and leads to embryonic lethality

To overcome the limitations of genetic loss-of-function approaches, we applied our previously optimised iLEXY system^34^, which induces nuclear export of a genetically-tagged protein in response to blue light. As the *spt5-HA-iLEXYs* line is homozygous viable and fertile when kept under safe light (“dark”), it allows both the maternally-deposited and zygotic SPT5 protein to be perturbed. We confirmed by immune-fluorescence that the SPT5-HA-iLEXYs fusion protein (hereafter referred to as SPT5) translocates from the nucleus to the cytoplasm upon blue light exposure (Fig. 3A). Importantly, depleting SPT5 for all of embryogenesis (mimicking a loss-of-function allele) results in complete embryonic lethality (**Fig. 3B**), in keeping with the conserved role of SPT5 as a universal elongation factor, and the lethality of a missense mutation (W049)^31^. This SPT5-HA-iLEXYs lethality is not caused by blue light toxicity, as wild-type embryos with the same blue light exposure are viable (**Fig. 3B**).

To quantify how efficiently SPT5 is depleted from chromatin throughout embryogenesis, we performed quantitative spike-in CUT&Tag after blue light exposure (using nuclei from the distant *Drosophila virilis* species as spike-in) at the beginning (3-4 h), middle (10-12 h) and end (18-20 h) of embryogenesis. The results show a global reduction in SPT5 binding to chromatin at all three time windows of 52%, 32%, and 37% at early, mid, and late stages of embryogenesis, respectively (**Fig. 3C, D**). These results indicate that iLEXY can reduce, but not abolish, SPT5 binding to chromatin throughout embryogenesis. The remaining level of SPT5 should prevent the degradation of RPB1 observed after strong SPT5 depletion in humans^24^, and therefore maintain Pol II binding (as we confirm below).

Taking advantage of the reversibility of the iLEXY system to dynamically modulate the nuclear levels of SPT5, we could precisely pinpoint the most sensitive embryonic time-windows to SPT5 depletion, which is impossible using classic loss-of-function approaches. Embryos were exposed to blue light for different time-windows, such that SPT5 was out of the nucleus and then brought back in (blue light ON→OFF) or in the nucleus and then depleted out (blue light OFF→ON) (**Fig. 3E**). This revealed an increased sensitivity to SPT5 depletion for embryonic viability starting from ∼12 h until 20 h, which roughly spans the second half of embryogenesis (**Fig. 3E**). To narrow down the sensitive time window even further, we exposed embryos to blue light at sliding 4-hour time intervals (**Fig. 3F**). In agreement with the previous experiment, embryonic lethality starts to occur (∼30%) when SPT5 is depleted between 8-12 h of embryogenesis, and reaches maximum levels (∼60%) between 12-16 h and 16-20 h (**Fig. 3F**). This indicates that the remaining level of SPT5 is sufficient for survival at early, but not at late embryonic stages, even though the depletion from chromatin is stronger at the early 3-4 h time window. Interestingly, compared to longer depletions (**Fig. 3E**), the lethality with a four-hour exposure is never complete and plateaus at ∼60% (**Fig. 3F**), suggesting that embryogenesis can recover to some extent from SPT5 depletion if it does not last too long.

To determine if there is recovery at the transcriptional level, we performed RNA-seq after depleting SPT5 for 4 h either between 8-12 h or between 12-16 h of embryogenesis, and then moved the embryos back to safelight conditions (where SPT5 can return to the nucleus) allowing the embryos to continue to develop (**Fig. 3G**). Depletion between 8-12 h results in the mis-expression of 92 genes compared to time-matched dark control embryos (|log_2_FC| > 1, adjusted p-value < 0.01), of which only eight are still misexpressed after 4 h of recovery, and one after 8 h (**Fig. 3G, Table S3**). Depleting SPT5 between 12-16 h leads to the misexpression of 224 genes. Remarkably, despite having a more severe effect on viability, within 4 h of being placed back in the dark the transcriptional profiles had almost completely recovered, with only 15 of these genes still misexpressed (**Fig. 3G, Table S3**). These results indicate that embryos can transcriptionally recover after SPT5 depletion, which also explains the incomplete lethality observed in the viability assays.

In summary, the iLEXY system facilitates nuclear depletion of SPT5 *in vivo* in living embryos, reducing – but not fully removing – SPT5 binding to chromatin to a similar extent across multiple stages of embryogenesis. While SPT5 is essential for embryogenesis, embryos are more sensitive to SPT5 depletion at later stages in the second half of embryogenesis, suggesting that SPT5 regulates different aspects of transcription at different embryonic stages– a hypothesis we explore in detail below. Even at later stages, embryos can transcriptionally recover if the depletion is not too prolonged, explaining the observed incomplete lethality of shorter exposures.

**Figure 3:**
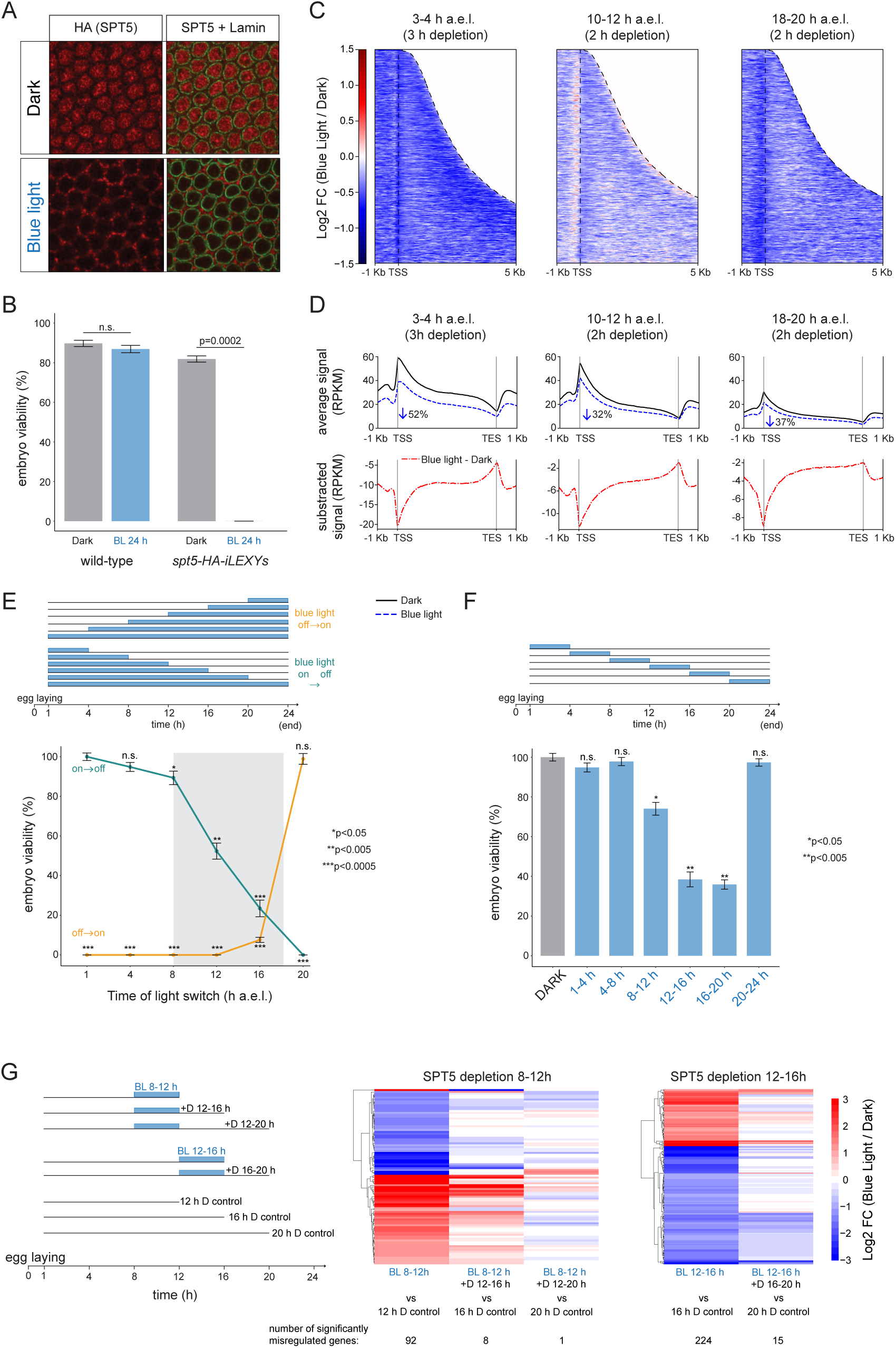
Optogenetic nuclear depletion of SPT5 decreases its binding to chromatin and leads to embryonic lethality. **(A)** Immunofluorescence of a stage-5 *spt5-HA-iLEXYs* embryo stained for SPT5 (HA tag, in red) and nuclear lamin (green) under safelight (Dark) or after 2 h of blue light exposure (lower). **(B)** Percentage embryonic viability (y-axis) comparing control and s*pt5-HA-iLEXYs* embryos in the dark (grey) and after blue light exposure for 24 h (BL), spanning all embryogenesis. Around 100 embryos were counted, in eac of the three independent replicates. Mean and standard deviations are shown, p-value from a two-sample t-test. **(C)** SPT5 depletion from chromatin using spike-in CUT&Tag with a SPT5 antibody at 3-4 h, 10-12 h and 18-20 h following a 3 or 2 h blue light exposure. Colours represent the quantitative log_2_ FC of SPT5 binding after blue light exposure relative to a time-matched dark control. Genes are ordered by increasing gene length, dashed lines (right) indicate the TES. Background signal beyond the TES was set to zero. For each timepoint only active genes (FPKM > 1) were considered – 3-4 h: N = 6581; 10-12 h: N = 8788; 18-20 h: N = 10008. **(D)** *Upper*: Metaprofiles of SPT5 binding for all genes following blue light induction (blue dashed line) compared to the dark control (black line) using genes from (C). *Lower*: Subtracted signal, blue light (BL) minus dark (D). Gene body is scaled from the TSS to the TES. The average depletion was calculated as the ratio between the TSS and TES signal in the dark and blue light (see Methods). **(E)** *Upper:* Embryo viability assays indicating the time windows of blue light exposure. Embryos were either shifted from dark (grey line) to blue light (blue rectangle), or vice versa. *Lower*: Mean viability percentages (y-axis), from 3 biological replicates (each with around 100 embryos), error bars represent standard deviations. Viability percentages for the dark condition were scaled to 100%, and all other data points adjusted accordingly. P-values from two-sample t-test comparing each depletion condition to its dark control. Shaded grey area indicates the time period with the highest sensitivity to SPT5 depletion. **(F)** *Upper:* Embryo viability assays following blue light exposure in 4-hour time windows across embryogenesis. *Lower*: As in (E). **(G)** *Left*: Schematic showing the time-window of blue light (BL) depletion (blue rectangle) followed by the indicated time window of recovery in the dark (D). SPT5 was depleted between 8-12 h and 12-16 h. *Right*: Log2 FC heatmaps of differentially expressed genes after blue light exposure compared to their respective dark control or recovery condition (indicated below the heatmaps). The number of significantly misregulated genes (|log2 FC| > 1 and adjusted p-value < 0.01) is indicated below each heatmap.

### SPT5 depletion during early embryogenesis causes a downstream shift of paused Pol II, defective elongation, and early transcriptional termination

Given our finding that different stages of embryogenesis are more or less sensitive to SPT5 depletion in terms of embryonic viability, we next investigated what aspect of transcription SPT5 regulates at early versus late stages of embryogenesis, starting with an early time point. A major, conserved event in early animal embryogenesis is zygotic genome activation (ZGA), during which transcription is initially established^44^. In *Drosophila*, this process occurs between 2-3 h of embryogenesis^45^ (**Fig. 4A**). To study the role of SPT5 in transcriptional regulation, we collected embryos just after this time-point, at 3-4 h, that had been exposed to blue light for either three hours (initiating depletion well before the onset of ZGA), or for one hour of blue light exposure (initiating during ZGA) (**Fig. 4A**). For each condition we examined changes in the occupancy of different components of the paused transcriptional machinery and in transcription after SPT5 depletion (**Fig. 4A**).

First, we examined SPT5 depletion from chromatin in the two conditions, which revealed a stronger global depletion after the longer 3 h exposure (**Fig. 4B**). While the 1 h depletion leads to a significant reduction in SPT5 signal at the gene body, there is little change in promoter-proximal binding. In contrast, the 3 h depletion reduces SPT5 binding at both the gene body and promoter-proximal region (**Fig. 4B**). This difference could be due to the longer depletion time or to the embryonic timing, as the longer depletion started well before the ZGA, *i.e*., before Pol II (and probably the transcriptional machinery) is assembled on chromatin at the majority of promoters^46^.

To examine the impact of SPT5 depletion on the paused transcriptional machinery, we performed spike-in CUT&Tag for the promoter-proximal and elongating Pol II (pSer5 and pSer2, respectively) and NELF. After 1 h depletion, the elongating Pol II pSer2 is more affected than the promoter-bound Pol II pSer5 (**Fig. 4C**), in keeping with the stronger reduction in SPT5 signal at the gene body compared to the promoter (**Fig. 4B**). Conversely, after 3 h – when SPT5 binding is also reduced at the promoter (**Fig. 4B**) – we detect stronger changes in Pol II binding at the promoter region (**Fig. 4C**). Overall, in both conditions, Pol II accumulates downstream of the TSS and then rapidly decreases until becoming comparable (if not mildly reduced) to the dark control within 0.5-1 kb from the TSS (**Fig. 4C**). Interestingly, the same happens with NELF (**Fig. 4C**). NELF normally binds to the promoter to stabilise pausing and once phosphorylated leaves the complex and therefore dissociates from chromatin^10,13^. However, after SPT5 depletion NELF unexpectedly remains associated with chromatin, moving downstream of the pausing site matching Pol II occupancy, suggesting that NELF and Pol II remain associated, where NELF is probably unphosphorylated.

To determine the extent of Pol II pausing and elongation after SPT5 depletion with higher resolution, we used PRO-seq to measure nascent RNA. After SPT5 depletion, the RNA peak corresponding to the pausing site is mildly reduced, while a second, prominent peak appears shortly downstream (**Fig. 4D**). To investigate this second peak, we compared the PRO-seq signal (nascent transcription) with nucleosome position, using MNase-seq data from early (ZGA) embryos^47^. In the dark control, the PRO-seq signal shows a main peak at +62 base pairs downstream of the TSS (pausing site) and a second, less obvious peak at +118, just before the +1 nucleosome (**Fig. 4E**). After SPT5 depletion, the PRO-seq signal is reduced at the canonical pausing site (+62) and increased at the +1 nucleosome (**Fig. 4E**), indicating that SPT5 depletion causes a downstream shift of Pol II, which gets stuck at the +1 nucleosome in agreement with it forming a second barrier to Pol II^48,49^. This was also observed in a human cell line after SPT5 depletion^24^, indicating that it is a conserved feature. Interestingly, weakly paused genes (bottom 25% from Fig. 2B) show no +62 peak but still have a +1 nucleosome peak, which increases after SPT5 depletion (**Fig. S3A, B**), indicating that even when pausing is absent at the canonical pausing site, SPT5 is required for Pol II to efficiently overcome the +1 nucleosome barrier.

Since SPT5 also acts as an elongation factor, we examined how SPT5 depletion affects elongation, both promoter-proximally and along the entire gene body. After SPT5 depletion, the PRO-seq signal rapidly decreases after the +1 nucleosome and becomes comparable to the dark control at ∼500 bp downstream of the TSS (**Fig. 4D, E**), indicative of early transcriptional termination around the first nucleosome. More downstream, between the TSS and TES, SPT5 depletion causes a gradual decay of nascent transcription along the gene body irrespective of the pausing strength associated with the promoter (**Fig. 4F**), which indicates defective elongation. Overall, SPT5 appears to both stabilise Pol II at the pausing site and to help Pol II to overcome the +1 nucleosome.

Finally, we assessed what effect this molecular phenotype has on the final RNA output. RNA-seq performed under the same conditions detected very minimal changes in total RNA abundance: three upregulated genes after 1 h of depletion, and five down- and four upregulated genes after 3 h of depletion (|log_2_FC| > 1, adjusted p-value < 0.01) (**Fig. 4G**). As RNA-seq measures steady state total RNA levels, any changes in zygotic transcription will likely be masked by the high maternal RNA load at this early embryonic stage, which explains why embryonic viability is not affected by SPT5 depletion at this early time point.

Taken together, our results indicate that in early embryogenesis the depletion of SPT5 causes a release of Pol II from the canonical pausing site to the more downstream +1 nucleosome where both Pol II and NELF accumulate. At the transcriptional level, this results in defective elongation along the gene body and early termination, in agreement with the dual role of SPT5 in pausing and elongation.

**Figure 4:**
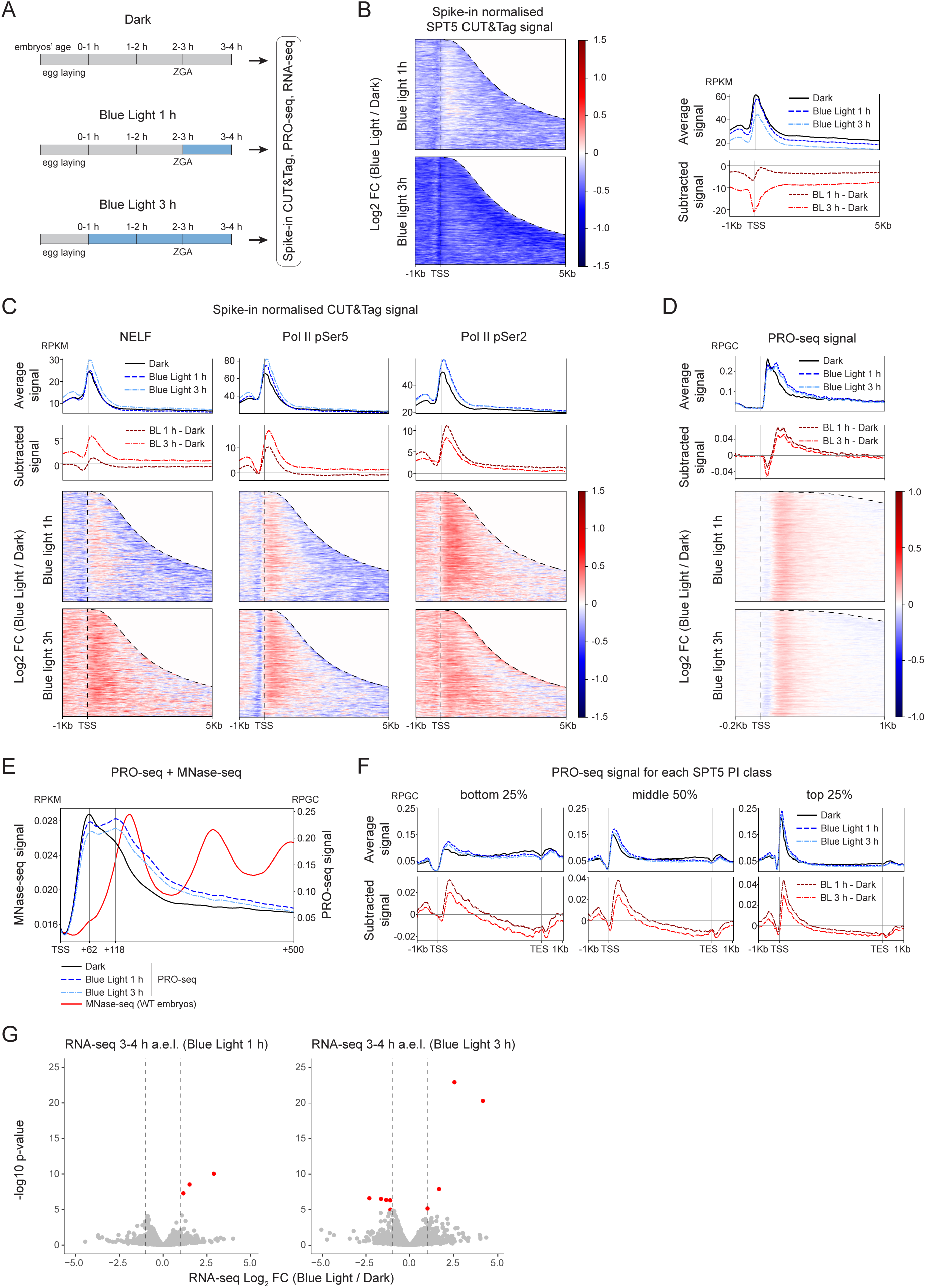
SPT5 depletion during early embryogenesis causes a downstream shift of paused Pol II, defective elongation, and early transcriptional termination. **(A)** Schematic showing the time window of SPT5 depletion (blue) in early embryogenesis used for spike-in CUT&Tag, PRO-seq, and RNA-seq. **(B)** SPT5 occupancy by spike-in CUT&Tag in 3-4 h embryos after blue light exposure for 1 or 3 hours, as depicted in (A). *Left*: Heatmaps of log_2_ FC in occupancy relative to dark controls for active genes. Genes (N = 6581) ordered by increasing gene length with the dashed line (right) indicating the TES. The signal beyond the TES was set to 0. *Right*: Metaprofiles of spike-in CUT&Tag signal in the dark and after 1 or 3 hours blue light depletion. Subtracted signal (red lines), blue light (BL) minus dark (D) shown below. **(C)** Same as (B), but for NELF, Pol II pSer5 and Pol II pSer2. Metaprofiles shown above the corresponding heatmaps. **(D)** PRO-seq in 3-4 h embryos after blue light exposure for 1 or 3 hours, as depicted in (A). *Upper*: Metaprofiles of PRO-seq signal for active genes in the dark and after 1 or 3 hours blue light depletion. Subtracted signal (red lines), blue light (BL) minus dark (D) shown below. *Lower*: heatmaps of log_2_ FC relative to dark controls. Genes ordered by increasing gene length with the dashed line (right) indicating the TES. For clarity, the signal beyond the TES was set to 0. **(E)** Metaprofiles of PRO-seq signal in the dark and after 1 or 3 hours blue light depletion, from the TSS to 500 bp downstream. A metaprofile of MNase-seq signal from early control embryos^47^ shown in red. The vertical dashed lines represent the position of the two detected peaks relative to the TSS, corresponding to the canonical pausing site and the +1 nucleosome. **(F)** Metaprofiles of PRO-seq signal for genes belonging to the bottom (N = 1392), middle (N = 2783), or top (N = 1392) SPT5 PI classes. The gene body is scaled from the TSS to the TES. **(G)** Volcano plots showing differentially expressed genes from RNA-seq at 3-4 h of embryogenesis after 1 or 3 hours blue light depletion compared to the dark control, as depicted in (A). Genes with significant changes (|log_2_ FC| > 1 and adjusted p-value < 0.01) are highlighted in red. Dashed vertical lines = |log_2_ FC| > 1.

### SPT5 depletion during late embryogenesis leads to pause release or defective elongation depending on the transcriptional and pausing state of the gene

To explore the role of SPT5 at later embryonic stages, we performed RNA-seq after depleting SPT5 at the same 4 h time windows when it leads to embryonic lethality (from **Fig. 3**), namely 8-12 h, 12-16 h, or 16-20 h, and the longer completely lethal window 8-20 h (**Fig. 5A**). At each time window, 48, 73, 187, and 523 genes are upregulated and 44, 151, 453, and 878 downregulated in their expression, respectively (|log_2_FC| > 1, adjusted p-value < 0.01) (**Fig. 5A**, **Table S5**). This progressive increase in the number of misregulated genes correlates with the decrease in viability, explaining the observed increase in embryonic lethality at later stages.

To characterise the molecular function of SPT5 during late embryonic stages, we performed spike-in CUT&Tag for SPT5, NELF and Pol II (pSer5 and pSer2) at 18-20 h after 2 h of SPT5 depletion. SPT5 occupancy is globally reduced, particularly at the promoter (**Fig. 5B**). In contrast to early embryogenesis, NELF does not move downstream of the pausing site together with Pol II (compare **Fig. 5B** to **4C**), but is instead mildly reduced. In contrast, Pol II (especially the elongating pSer2) appears to accumulate downstream of the TSS at the beginning of the gene body (**Fig. 5B**). PRO-seq revealed a striking difference in the impact on nascent transcription compared to early embryogenesis (compare **Fig. 5C, D** and **4D, E**). While SPT5 depletion causes a decrease in pausing at the canonical pausing site also at 18-20 h, the released Pol II does not stall at the +1 nucleosome at this late stage (**Fig. 5C, 5D**), independent of the level of pausing (**Fig. S4A, B**). The comparison between the dark control at 3-4 h versus 18-20 h shows that in late embryogenesis Pol II pauses only at the canonical pausing site (**Fig. S4C**), suggesting that the +1 nucleosome is not a strong barrier for Pol II progression at late embryonic stages, in contrast to the early time points.

To investigate how changes in the occupancy of the transcriptional machinery are related to transcriptional changes (**Fig. 5A**), we generated metaprofiles of the CUT&Tag signal in control (dark) and SPT5 depleted embryos, for genes that were upregulated, downregulated or unchanged in their expression after SPT5 depletion between 16-20 h of embryogenesis (from **Fig. 5A**). Upregulated genes have increased Pol II pSer5 signal close to their promoter and increased Pol II pSer2 signal all the way from the TSS to the TES (**Fig. 5E**). This indicates that at this late time point – unlike early embryogenesis – the transcriptional machinery released from the promoter can transcribe until the TES and increase productive transcription, suggestive of pause release. Interestingly, NELF does not remain associated with the released Pol II and its promoter occupancy is reduced (**Fig. S4D**), again in contrast to early stages. On the other hand, downregulated genes after SPT5 depletion have mildly reduced Pol II binding along their gene body (**Fig. 5E**), suggestive of impaired elongation. Downregulated genes also have less accumulation of Pol II at their promoters under control (dark) conditions compared to upregulated genes, suggesting that they are inherently less regulated by pausing (**Fig. 5E**). This is supported by their SPT5 PI, which is significantly lower for down-compared to up-regulated genes (**Fig. S4E**). This suggests that upregulation is caused by pause release while downregulation is due to defective elongation. To test this hypothesis, we plotted the average PRO-seq signal for up- and downregulated genes (**Fig. 5F**). For upregulated genes, nascent transcription is increased all the way from the TSS to the TES, in contrast to genes that do not change, where the signal is only slightly increased just downstream of the pausing site, but becomes comparable to the dark control towards the TES (**Fig. 5F**, upper). Interestingly, upregulated genes also show a higher PRO-seq peak at their TSS (**Fig. 5F**). As pausing can prevent new initiation^42^, this suggests that the decrease in pausing for upregulated genes is accompanied by an increase in initiation. At downregulated genes – which do not show promoter proximal pausing – the PRO-seq signal is reduced along the gene body (**Fig. 5F**). Taken together, these data indicate that SPT5 depletion can lead to both upregulation (through pause release into productive elongation) and downregulation (through impaired elongation) of gene expression, depending on a gene’s pausing level.

Next, we asked whether the transcriptional dynamics of a gene, in addition to its pausing state, can influence the transcriptional response to SPT5 depletion. To assess this, we took advantage of the highly dynamic changes in gene expression during embryogenesis. For each depletion time window from Fig. 5A, we used a wild-type time-course of RNA-seq^35^ to classify the misregulated genes as physiologically increasing or decreasing in their expression over the embryonic time window of depletion (log_2_[late/early] > 1 or < -1, respectively) (**Fig. 5G**). Of the 878 downregulated genes after SPT5 depletion between 8-20 h (**Fig. 5A**), 746 (85%) normally increase in their expression during this time window, while only 47 (5%) decrease (**Fig. 5G**). Conversely, of the 523 upregulated genes after SPT5 depletion, 338 (65%) are normally decreasing in their expression at these stages of embryogenesis, while 145 (28%) increase (**Fig. 5G**). The same trend holds true across different embryonic time windows, especially for the downregulated genes (**Fig. S4F**). This indicates that SPT5 depletion impacts genes with dynamic expression in opposite ways, causing the downregulation of genes that are increasing in expression (and likely more susceptible to changes in elongation) and the upregulation of those decreasing in expression. This is in agreement with our finding that pausing both precedes and follows active transcription (**Fig. 2E**), and supports the hypothesis that SPT5 and pausing are important both to begin and end active transcription.

In summary, our results indicate that SPT5 depletion in late embryogenesis causes the release of paused Pol II from the canonical pausing site into the gene body, without getting stuck at the +1 nucleosome. This leads to both up- and downregulation of gene expression depending both on the level of pausing and the transcriptional dynamics of the gene. Paused genes can undergo pause release and result in upregulation, while non paused genes are more susceptible to a block in elongation leading to their downregulation. This is exacerbated for genes that are physiologically ramping up in their expression, and therefore their effective elongation. In this way, genes that are normally increasing or decreasing in expression are more prone to be downregulated or upregulated, respectively, indicating that SPT5 contributes to fine-tuning dynamic changes in gene expression.

**Figure 5:**
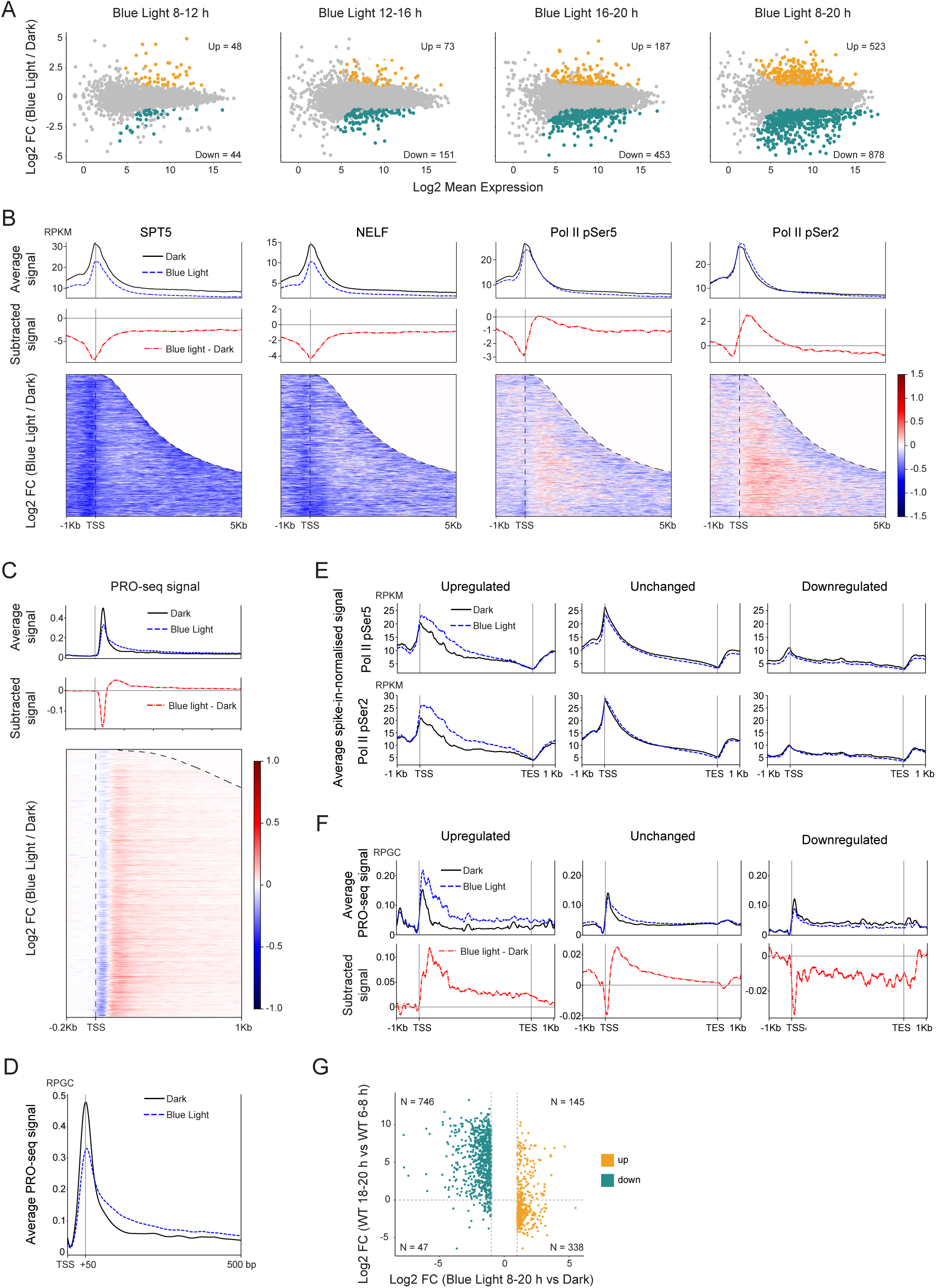
SPT5 depletion during late embryogenesis leads to pause release or defective elongation depending on the transcriptional and pausing state of the gene. **(A)** Mean average (MA) plots showing differentially expressed genes after four hours of blue light exposure at different embryonic time windows: 8-12, 12-16, and 16-20 h, and 12 hours depletion, from 8-20 h. Genes with significantly upregulated or downregulated expression are indicated (|log_2_ FC| > 1 and adjusted p-value < 0.01). **(B)** SPT5, NELF, Pol II pSer2, and Pol II pSer5 occupancy at 18-20 h after 2 h blue light exposure. *Upper*: metaprofiles of CUT&Tag signal for active genes (FPKM > 1, N = 10008) for each factor in the dark (black lines) and after 2 h blue light exposure (blue dashed lines). Subtracted signal (red dashed line), blue light (BL) minus dark (D) shown below. Displayed from 1 kb upstream to 5 kb downstream of the TSS (vertical dashed line), binned in 10 bp bins. *Below*: heatmaps of log₂ FC in binding for each factor relative to its dark control, shown from 1 kb upstream to 5 kb downstream of the TSS. Genes ordered by gene length, from the shortest (top) to the longest (bottom), with dashed lines (right) indicating the TES. For clarity, signal values beyond the TES were set to zero. **(C)** PRO-seq in 18-20 h embryos after blue light exposure for 2 h. *Upper*: Average signal for active genes (FPKM > 1, N = 10008) in the dark and after 2 h blue light exposure. Subtracted signal (red lines), blue light (BL) minus dark (D) shown below, displayed from 0.2 kb upstream to 1 kb downstream of the TSS. *Lower*: heatmap of log_2_ FC relative to dark controls. Genes ordered by increasing gene length, with the dashed line (right) indicating the TES. The signal beyond the TES was set to 0. **(D)** Zoom-in of the TSS region showing average PRO-seq signal from (C). The vertical dashed line corresponds to the canonical pausing site. **(E, F)** Metaprofiles of Pol II pSer5 and pSer2 (**E**) and PRO-seq (**F**) signal on genes upregulated, downregulated or unchanged after SPT5 depletion at 16-20 h (from panel (A)). **(G)** Scatter plot showing the relationship between the changes in gene expression after SPT5 depletion (x-axis, log₂ fold blue light vs dark at 8-20 h) and genes with physiological changes in their expression during embryogenesis from 6-8 h to 18-20 h (y-axis, log₂ ratio of expression). Genes above 0 on the y-axis (horizontal dashed line) increase in expression over developmental time in wild-type embryos, while the ones below 0 decrease. Significantly upregulated or downregulated genes from (A), which are expressed at 6-8 h and 18-20 h (FPKM > 1), are shown.

## DISCUSSION

### Pol II pausing throughout embryogenesis

Examining how Pol II pausing changes over the course of embryogenesis, our results indicate that SPT5 promoter occupancy (and pausing) is established before transcription starts, reminiscent of poised promoters. The onset of active transcription is accompanied by pause release into productive elongation, which is sometimes accompanied by a higher rate of initiation. Depending on whether pause release is higher or lower than the rate of new initiation, this will result in a decrease or increase in the pausing index, respectively – even if elongation (and thus the transcriptional output) increases similarly over time for the two groups of genes. Interestingly, when transcription ends (i.e., when genes are switching off), there is also an accumulation of SPT5 at the promoter and thus a gain in pausing for many genes. Pausing therefore changes not only in relation to active expression, but also to its onset and end, suggesting that it plays a widespread role in the dynamics of gene expression.

Promoters of highly paused genes have high SPT5/Pol II promoter accumulation and are bound by NELF. In contrast, genes with low pausing show little or no SPT5/Pol II promoter accumulation and are generally not bound by NELF. Interestingly, the elongation factors SPT6 and PAF1 bind to the same protein binding pocket on Pol II as NELF^13,14^. Our data therefore supports a model where the promoters of weakly paused genes have no NELF, leaving Pol II readily accessible to the elongation factors SPT6 and PAF1. Conversely, at the promoter of highly paused genes, SPT5/Pol II slow down and are stabilised by NELF binding. When NELF is bound, Pol II is no longer accessible to the elongation factors until NELF becomes phosphorylated and dissociates from the complex. This property of a promoter to be highly paused, and accumulate SPT5/Pol II binding, is constant over time, as promoters are generally either bound or not bound by NELF across the whole of embryogenesis, suggesting that this is promoter specific and likely encoded in the promoter’s sequence. In agreement, highly paused promoters are enriched in specific motifs^3,43^ whose causal role in pausing has been assessed for specific loci, for example by mutating promoter motifs^39^ or by substituting a highly paused with a weakly paused promoter – which resulted in less synchronous gene expression, with more variability across a field of cells during gastrulation^38^. Finally, we find an intriguing link between the extent of pausing and gene length: highly paused genes are on average five times longer than weakly paused genes (Fig. S2F). While the advantage that this might impart on the transcription of long genes remains unclear, it hints at a selection of the promoter type, again suggesting that pausing is genetically encoded within the promoter sequence.

### SPT5 modulates different aspects of transcription in early versus late embryogenesis

Depleting SPT5 at specific time windows uncovered different sensitives to SPT5 depletion at different stages of embryogenesis (**Fig. 6A**). This very unexpected finding suggests that SPT5 has different molecular functions at early versus late embryonic stages. By measuring NEFL and Pol II occupancy and transcription, our results show that SPT5 depletion in the early embryo results in a downstream shift of Pol II from the pausing site to the +1 nucleosome. Pol II is not able to efficiently overcome the +1 nucleosome barrier and nascent transcription (as seen by PRO-seq) is reduced, compatible with early termination. SPT5 depletion also results in a reduction of Pol II signal and transcription to the end of the gene, indicating defective elongation. Interestingly, NELF also moves downstream with Pol II, which has not been observed before. Since NELF is unphosphorylated when bound to the promoter and dissociates once phosphorylated, it is likely that NELF is in an unphosphorylated state when it shifts downstream together with Pol II. Our results therefore show two molecular phenotypes after SPT5 depletion in early embryos – an inability of Pol II to transverse the +1 nucleosome and an aberrant association between NELF with the Pol II machinery, which both likely contribute to early transcriptional termination. At this early stage, this does not lead to changes in total RNA abundance, and thus to embryonic lethality (**Fig. 6A**, left).

During late embryogenesis, the depletion of SPT5 does result in transcriptional changes, which correlate with developmental time: the later the developmental stage, the larger the number of misregulated genes. SPT5 depletion leads to a reduction in NELF binding, as also observed in mammalian cells^24^, with no movement into the gene body, in contrast to early stages. Perhaps it is this lack of the aberrant association between NELF and Pol II in the gene body that enables productive transcription. Another key difference from early embryonic stages, is the ability of Pol II to enter the gene body (seen by CUT&Tag) and transcribe (PRO-seq) past the +1 nucleosome at 18-20 h, indicating that Pol II can overcome this barrier at late stages, resulting in either the up- or downregulation of gene expression (**Fig. 6B**, right).

The barrier of the +1 nucleosome for Pol II elongation seems to be a key difference between early and late stages of embryogenesis. This is actually already visible in the control (non-depleted) embryos. While at 18-20 h there is only one PRO-seq peak corresponding to the pausing site, at 3-4 h there is a second peak at the +1 nucleosome. This interesting feature might be a residual effect of ZGA, which mostly happens at nuclear cycle 14, between 2-3 h. One theory for how the zygotic genome is activated is through histone dilution – the mother supplies a huge amount of histone proteins into the egg, leading to a very high nucleosome concentration. During each of the 13 successive cell divisions before transcription, the nucleosome concentration becomes progressively diluted, leading to more spacing between nucleosomes, making chromatin more permissive for transcription^44^. In agreement, the +1 nucleosome was shown to play a role in ZGA in *Drosophila* embryos^47^.

### The transcriptional response to SPT5 depletion in late embryogenesis depends on the gene’s level of pausing and transcriptional state

The dynamic changes in gene expression during late embryogenesis uncovered an interesting feature – SPT5 depletion can lead to both an up- and downregulation of gene expression (**Fig. 6B**). Upregulated genes have higher Pol II occupancy and nascent transcription along their gene body indicating more Pol II released from the promoter, which can efficiently travel all the way to the TES. Since in human cell lines the degradation pathway is activated on chromatin^24^, the fact that Pol II occupancy is higher for these genes demonstrates that Pol II is not degraded or disassociated from chromatin after partial SPT5 depletion in our system. Under control conditions (dark), upregulated genes have a high SPT5/Pol II and PRO-seq signal at their promoter and are bound by NELF, all pointing to strong pausing. After SPT5 depletion, they have even higher levels of Pol II at their TSS, in agreement with findings that pausing inhibits new initiation^42^ and suggests that SPT5 depletion, by decreasing pausing, might allow new initiation (**Fig. 6B**, left). In contrast, downregulated genes in control conditions typically have little SPT5/Pol II and PRO-seq promoter signal, and are not bound by NELF, indicating less regulation by pausing (**Fig. 6B**, right). This favours a model where SPT5 depletion from genes that are regulated by pausing leads to their upregulation, while from genes that mostly rely on elongation results in their downregulation (**Fig. 6B**).

The response of genes to SPT5 depletion – whether it is up- or downregulated – also depends on the genes’ dynamics in terms of their developmental regulation. Genes that are physiologically increasing in their expression require an increase in Pol II elongation. In this context, SPT5 depletion leads to a downregulation in their expression. Genes that are normally decreasing in their expression have less Pol II elongation and likely more pausing. In this context, SPT5 depletion, as it releases pausing, can lead to an upregulation in expression. The transcriptional response to SPT5 depletion will therefore depend on two properties of the gene – its inherent level of pausing (regulated by sequence features in its promoter) and the dynamic regulation of the gene’s expression, whether it is being switched on or off. SPT5 can therefore fine-tune transcription through different mechanisms during embryogenesis.

**Figure 6:**
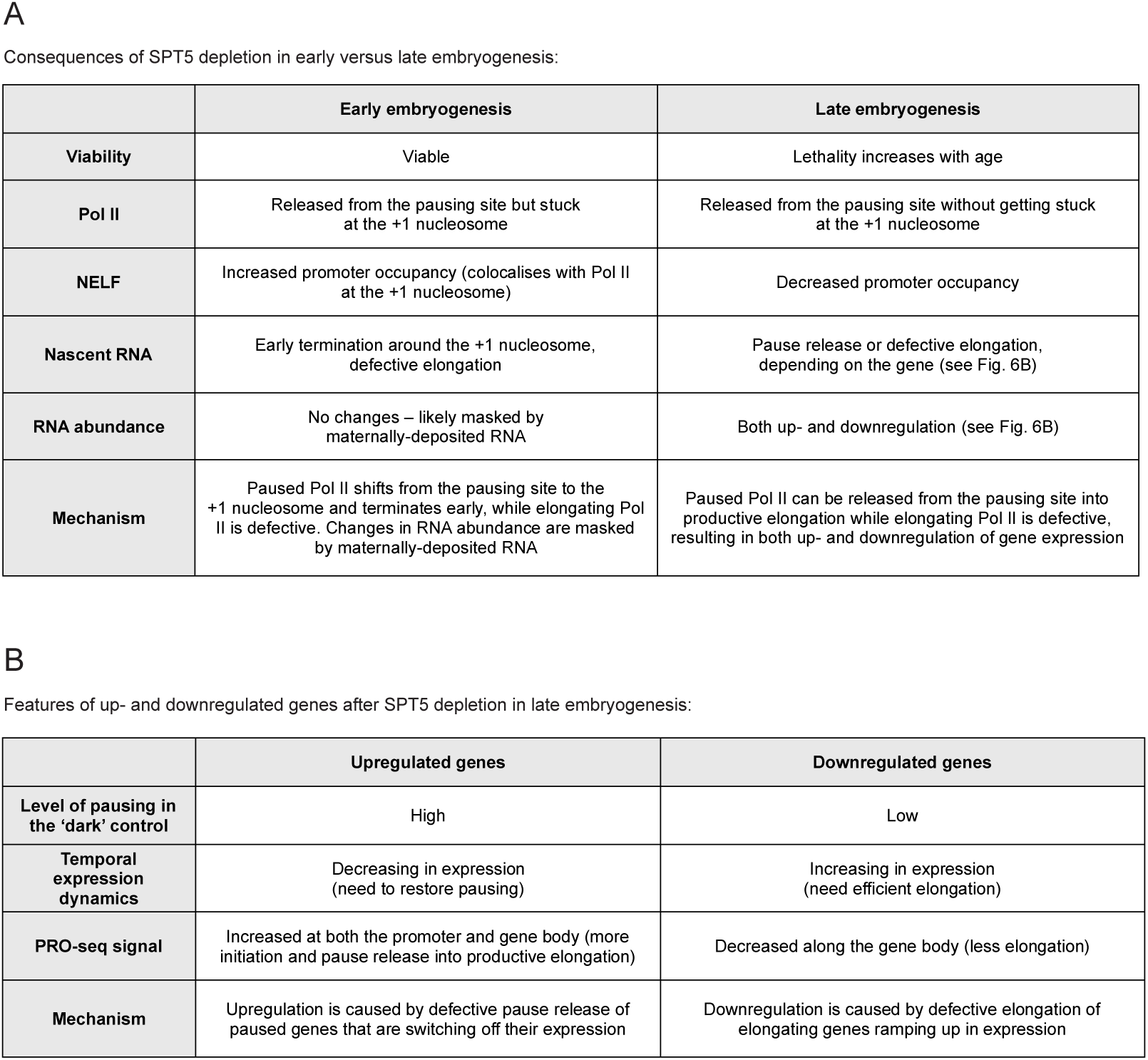
Summary of the phenotypes observed after SPT5 depletion during embryogenesis. **(A)** Overview of the embryonic and molecular phenotypes observed in early (left) and late (right) stages of embryogenesis **(B)** Molecular features of up (left) and down (right) regulated genes after SPT5 depletion at late stages of embryogenesis

### Limitations of the study

Although the incomplete depletion of SPT5 is desirable to avoid the degradation of RBP1, a 100% depletion (while somehow also blocking Pol II degradation in embryos) would be interesting to assess the role of Pol II pausing for early stages of embryogenesis. Our results show that SPT5 depletion leads to molecular phenotypes at both early and late stages of embryogenesis, but only causes embryonic viability phenotypes at late stages. It is interesting to speculate why early embryonic stages are less sensitive to SPT5 depletion in terms of embryonic viability. Does this suggest that Pol II pausing is not essential at early stages of embryogenesis? We don’t think so – rather the opposite. Our results reveal very high levels of SPT5-Pol II binding to the promoter at early stages, such that depleting the complex by 50% may still leave high enough absolute protein levels to mediate Pol II pausing. A key interesting difference between early and late stages is the ability of Pol II to move past the +1 nucleosome at the transition between pausing and elongation. Perhaps this second regulatory step, the +1 nucleosome, is required because pausing is so important during early embryogenesis. Future work is needed to address this interplay between the two (pausing site and +1 nucleosome), but this will be very challenging in embryos, as it requires depleting SPT5 and perturbing pausing at the +1 nucleosome.

## RESOURCE AVAILABILITY

### Lead contact

Further information and requests for resources and reagents should be directed to and will be fulfilled by the lead contact Eileen Furlong.

### Materials Availability

The *spt5-HA-iLEXYs* fly line used in this study was generated by the authors as described in the methods section and is maintained for the community by the Furlong lab at EMBL. Only costs to cover shipping and packaging will be requested.

### Data and code availability

- All raw data has been deposited to EMBL-EBI ENA public repository (https://www.ebi.ac.uk/arrayexpress/browse.html). The accession numbers are listed in the key resources table. The processed bigWigs of CUT&Tag data for SPT5, NELF, Pol II pSer5 and pSer2, and RNA-seq and PRO-seq data in dark and blue light depleted SPT5-iLEXY embryos at multiple stages are available on the Furlong lab web page, http://furlonglab.embl.de/data
- The code required to reproduce the figures is available at https://git.embl.de/grp-furlong/shared/spt5-ilexy
- Any additional information required to reanalyse the data reported in this paper is available from the lead contact upon request

## ACKNOWLEDGEMENTS

The authors thank all Furlong lab members for helpful comments. We are very grateful to John Lis for the guinea pig anti-SPT5 antibody and Julia Zeitlinger for the rabbit anti-NELF antibody. This work was technically supported by EMBL’s Genomics, Protein Expression and Purification, Advanced Light Microscopy and *Drosophila* injection Core facilities, and by the external resources Bloomington *Drosophila* Stock center and Flybase. We are very grateful for financial support from Deutsche Forschungsgemeinschaft (DFG SPP 2202) and European Research Council (ERC advanced grant) agreement 787611 (DeCRyPT) grants awarded to E.E.M.F.

## AUTHOR CONTRIBUTIONS

A.D. and E.E.M.F conceptualised, designed and planned the study. A.D. performed all experiments, with help from A.G. for PRO-seq and R.R.V. for nuclei extractions. M.M. and N.E. performed data analyses with input and support from M.F. Y.K. and S.F. designed and generated the *spt5-HA-iLEXYs* knock-in line. A.D. and E.E.M.F. interpreted the results and wrote the manuscript, with input from all authors. E.E.M.F. supervised the project and provided funding.

## STAR METHODS

### EXPERIMENTAL MODEL AND STUDY PARTICIPANT DETAILS

*Drosophila melanogaster* (*D. melanogaster*) and *Drosophila virilis* (*D. virilis*) stocks were maintained in bottles containing a standard cornmeal-agar medium and grown at 18°C or at 25°C. To generate the CRISPR-Cas9 mediated knock-in of the iLEXYs tag^34^ into the *spt5* endogenous locus, we used the line w[1118]; Pbac{y[+mDint2] = vas-Cas9}VK00027 (*vasa-Cas9*) (Bloomington stock 51324). The iLEXY lines were always maintained and handled under safelight conditions, covering all light sources with amber light filter foil.

### METHOD DETAILS

#### Generation of the s*pt5-HA-iLEXYs* line

To tag the endogenous *spt5* gene with the iLEXYs cassette^34^, we used the scarless CRISPR-Cas9 gene editing technology (https:// flycrispr.org/scarless-gene-editing/). The flyCRISPR Target Finder programme (https://flycrispr.org/target-finder/) was used to design gRNAs close to the *spt5* stop codon. The annealed oligos gRNA_FW: 5’-CTTCGTATGTAGAGCTTAGTCGA-3’ and gRNA_REV: 5’-AAACTCGACTAAGCTCTACATAC-3’ were inserted into the BbsI site of the pU6-BbsI-gRNA vector^52^. A fusion product of the *spt5* left homology arm (1179 bp directly upstream of the stop codon, amplified from the targeted fly line), the HA tag, the iLEXY module, and the genomic region between the gRNA cleavage site and the TTAA sequence were inserted into the AarI site of the pHD-DsRed-Scarless donor vector^52^. The *spt5* right homology arm (spanning the TTAA and the 1150 bp downstream) was amplified from the targeted fly line and cloned into the SapI site of the same plasmid.

CRISPR donor and gRNA plasmids were combined at a 2:1 molar ratio and injected into *vasa-Cas9* embryos (Bloomington stock 51324) by EMBL’s *Drosophila* injection service. Positive transformants among F1 flies were identified by DsRed expression in the eyes and used to establish stable homozygous stocks. The knock-in was confirmed by PCR amplification and subsequent Sanger sequencing of the CRISPR allele, from outside of the left homology arm to outside of the right homology arm^52^.

#### Blue light induced nuclear depletion

Blue light illumination of *D. melanogaster* embryos was performed using an updated version of the previously described^34^ custom-made, programmable LED-based blue light illumination box. The illumination was performed alternating 2-second blue light pulses at 30% intensity with 1-second breaks, at 25°C.

#### Embryo viability assays

Embryo viability was quantified by determining the hatching rate of embryos into first-instar larvae. After three 1-hour pre-lays, embryos were collected for one hour at 25°C on apple juice agar plates, the yeast paste was removed, and the embryos were counted (the light of the stereoscope was covered with amber filter foil). The plates were kept at 25°C under safelight conditions and only exposed to the blue light in the LED box (also at 25°C) at the desired time window, as described for each experiment. Thirty hours after egg laying, the number of hatched and unhatched eggs were counted to determine the hatching rate. Each condition was tested in triplicate, using around 100 embryos per replicate.

#### Immunostaining

Dechorionated embryos were fixed in 4% methanol-free ultra-pure formaldehyde (Polysciences #18814-20) in PBS for 15 min and stored in methanol at -20°C as required. Before use, embryos were rehydrated in PBT (PBS with 0.2% Triton X-100) and blocked in PBT-BSA (PBT with 10% BSA) for 1 h. Embryos were incubated overnight with the primary rabbit anti-HA antibody (Cell Signaling #3724) and mouse anti-Lamin antibody (DSHB #ADL101), both diluted 1:250 in PBT-BSA, followed by five washes in PBT and incubated for 1 h with a secondary biotinylated anti-rabbit antibody (VectorLabs #BA-1000), diluted 1:500. After five PBT washes, they were incubated with Streptavidin, Rhodamine Red™-X conjugate (ThermoFisher Scientific #S6366) and the Goat anti-Mouse IgG1 Cross-Adsorbed Secondary Antibody, Alexa Fluor™ 647 (ThermoFisher Scientific #A21240), both diluted 1:500, for 1 h. Embryos were mounted in ProLong^TM^ Gold with DAPI (ThermoFisher Scientific #P36931) and imaged on a Zeiss LSM 780 using the 63x/1.4 Oil DIC M27 objective.

#### RNA-seq

Dechorionated embryos were snap-frozen in liquid nitrogen and stored at -80°C. Embryos were homogenised in TRIzol LS (ThermoFisher Scientific #10296028) with a pestle on ice. RNA was isolated using standard phenol-chloroform extraction, and the remaining DNA digested with RNase-free DNase I (Roche #4716728001) for 30 min. RNA quality was checked on a Bioanalyzer (Agilent). For the depletion experiments at 3-4 h (Fig. 3), RNA-seq was performed using the Ovation RNA-Seq System 1-16 for Model Organisms (from NuGEN, now Tecan Genomics) following the manufacturer’s protocol. For all other experiments (Fig. 3 and Fig. 5), RNA-seq was performed from 1 μg of total RNA using the NEBNext Ultra Directional RNA Library Prep Kit for Illumina (NEB #E7420) with the NEBNext Poly(A) mRNA Magnetic Isolation Module (NEB #E7490) according to the manufacturer’s instructions. All RNA-seq experiments were performed with two biological replicates. Final libraries were run on a Bioanalyzer, multiplexed and sequenced on Illumina platform at the EMBL Genomics Core Facility.

#### PRO-seq

Dechorionated embryos were snap-frozen in liquid nitrogen and stored at -80°C. Quick precision run-on sequencing (qPRO-seq) was performed as previously described^29^ with minor modifications. Commercial RNase inhibitors were replaced by recombinant porcine MBP-RNasin (EMBL PEP core facility), at a final concentration of 10 µg/mL. Approximately 50 mg of frozen dechorionated embryos were used for nuclei extraction for each replicate. For 18-20 h embryos, five additional strokes with the loose pestle were necessary for proper homogenisation. Between 2-4 million nuclei were used in each run-on reaction. Final libraries were amplified from 5 µL of reverse transcription product, with a total of 20 PCR cycles. The final libraries were quantified with Qubit, run on the Bioanalyzer with HS DNA reagents, and sequenced at the EMBL Genomics Core Facility.

#### CUT&Tag and spike-in CUT&Tag

***Embryo collections:*** Embryos at the appropriate stage were dechorionated, and fixed in 1.8% formaldehyde (Sigma-Aldrich #1.04003.1000) in cross-linking solution (50 mM Hepes, 1 mM EDTA, 0.5 mM EGTA, 100 mM NaCl, pH 8) for 15 min. Formaldehyde was quenched with 125 mM Glycine (Millipore #1.04201.1000) in PBT (PBS with 0.1% Triton-X100). After two washes in PBT, embryos were dried on a Nitex membrane (Sefar #03-125/45), snap-frozen in liquid nitrogen and stored at -80°C until further use.

***Nuclei extraction:*** Nuclei from *D. melanogaster* and *D. virilis* embryos were isolated as previously described^53^. After isolation, nuclei were counted with DAPI staining using a FACSymphony^TM^ A3 cell analyser together with CountBright Absolute Counting Beads (Thermo scientific C36950) at the EMBL Flow Cytometry Facility. Nuclei were resuspended in nuclear freezing buffer (50 mM Tris pH 8, 25% glycerol, 5 mM MgAc2, 0.1mM EDTA, 5 mM DTT), aliquoted, snap-frozen in liquid nitrogen, and stored at -80°C until further use.

***Nuclei sorting:*** For the timecourse of SPT5 occupancy on the *spt5-HA-iLEXYs* line (using an anti-HA antibody), the use of a spike-in was not possible, as the anti-HA antibody would not recognise any epitope in the *D. virilis* spike-in. Therefore, to ensure that an equal amount of input material was used, we sorted 100,000 single nuclei from each time point, in duplicate, using the BD FACSAria^TM^ Fusion. For all the other CUT&Tag experiments, a spike-in was used (spike-in CUT&Tag). To ensure the correct balancing of the spike-in, single nuclei were sorted using the BD FACSAria^TM^ Fusion. Briefly, 50,000 *D. melanogaster* nuclei were sorted together with 50,000 *D. virilis* nuclei into 1 mL of ice-cold dig-wash buffer (20 mM HEPES pH 7.5, 150 mM NaCl, 0.5 mM Spermidine, 0.01% Digitonin, 1x Roche cOmplete Protease inhibitors) and kept on ice until all sorted samples were ready.

***Antibody/pA-Tn5 binding, tagmentation, and library preparation:*** CUT&Tag was performed as previously described^54^. Briefly, sorted nuclei were bound to Concanavalin A beads (Polysciences Europe GmbH, #86057-3) and incubated with primary antibody, secondary antibody, and finally with pA-Tn5 (EMBL PEP core facility). For tagmentation, nuclei were resuspended in 200 μL of CUT&Tag tagmentation buffer (36.3 mM Tris-Acetate pH 7.8, 72.6 mM K-Acetate, 11 mM Mg-Acetate, 17.6% Dimethylformamide [Sigma #D4551]) and incubated at 37°C for 1 h. The reaction was stopped and samples were de-crosslinked overnight (25 mM Tris pH 8, 50 mM EDTA, 0.1% SDS, 50 mM NaCl, 0.5 mg/ml Proteinase K [Qiagen #19131]). The next day, DNA was purified by column purification (DNA clean and Concentator-5 Kit, Zymo #D4014) and treated with RNase. The PCR for library preparation was performed using P5 and P7 primers using the NEBNext® High-Fidelity 2x PCR Mater Mix (M0541L). The PCR products were purified with 1.3x Agencourt AMPure XP beads (Beckman Coulter, #A63881), quantified with Qubit, run on the Bioanalyzer with HS DNA reagents, and sequenced.

The following primary antibodies were used: rabbit anti-HA (Cell Signaling #3724), guinea pig anti-SPT5c^55^ (gift from John Lis), rabbit anti-NELF-E (gift from Julia Zeitlinger), mouse anti-Pol II pSer5 (Antibodies-online #ABIN6655367), and mouse anti-Pol II pSer2 (Antibodies-online #ABIN6655366). The secondary antibodies were: guinea pig anti-rabbit IgG (Antibodies-online #ABIN6923140), goat anti-guinea pig IgG (Antibodies-online, ABIN6923140), and rabbit anti-mouse IgG (AbCam #ab46540).

### QUANTIFICATION AND STATISTICAL ANALYSIS

#### RNA-seq data analysis

***QC and coverage tracks:*** FastQC^56^ (v. 0.11.8) was used to check sequencing quality. Reads were trimmed with TrimGalore^57^ (v. 0.6.7) requiring a minimum base quality of 20 and a minimum adapter overlap of 2. Trimmed FASTQ files were mapped to the dm6 reference genome using STAR^58^ (v. 2.7.9) with the options ‘--readFilesCommand zcat -- outSAMtype BAM Unsorted --outSAMunmapped Within’. Problematic SAM records were filtered using Picard’s CleanSam^59^ (v. 2.16.0) and the resulting BAM file was sorted and indexed with SAMtools^60^ (v. 1.9). PCR duplicates were identified using Picard MarkDuplicates (v. 2.16.0) and the options ‘CREATE_INDEX=true REMOVE_DUPLICATES=false ASSUME_SORT_ORDER=coordinate SORTING_COLLECTION_SIZE_RATIO=0.25’. To check mapping quality of the reads to the *D. melanogaster* genome, FastQ Screen^61^ (v. 0.15.2) was used with a subset of 1000000 reads. Different quality control metrics were obtained by running Picard CollectRnaSeqMetrics (v. 2.16.0) using the refflat file for gene annotation, a list of ribosomal intervals for the *Drosophila melanogaster* reference genome version 6.13^62^ and a rRNA fragment percentage of 0.001. Insert size distribution was assessed with Picard CollectInsertSizeMetrics (v. 2.16.0). Correlation between samples was checked with deepTools^63^ multiBamSummary and plotCorrelation (v. 3.5.0). Signal tracks (bigwig) were obtained using deepTools bamCoverage (v. 3.5.0) with options ‘--binSize 10 -- effectiveGenomeSize 125464728 --extendReads --exactScaling --ignoreDuplicates -- normalizeUsing ’RPKM’ --ignoreForNormalization chrX chrM’. Coverage for replicate was then averaged with deepTools bigwigCompare (v. 3.5.0) with options ‘--operation mean -- binSize 10’. QC results were summarized and compared across samples with multiQC^64^ (v. 1.11).

***Differential expression analysis:*** The read alignment and gene expression quantification was performed with the RSEM^65^ pipeline (v. 1.3.3) with options “--paired-end --star --no-bam-output --star-gzipped-read-file” and based on FlyBase^66^ release 6.37 gene annotation, with the BDGP6.46 genome assembly. For the RSEM pipeline the option ‘--strandedness’ was set to ‘forward’ for RNA-seq data from the depletions at 3-4 h and to ‘reverse’ for all other experiments (see “RNA-seq” experimental methods section). RSEM expected counts were rounded to the closest integer and used as input for DESeq2^67^ (v. 1.38.3). Metadata and contrast files were provided as input, along with normalized gene expression tables generated by RSEM. Genes with a FPKM greater than 1 in each condition (*e.g.*, *SPT5-iLEXYs* dark [two replicates] and blue light [two replicates] samples) were considered expressed and tested for differential expression. Differentially expressed genes were defined as |log_2_FC| > 1 and FDR < 0.01.

#### CUT&Tag data analysis (timecourse of SPT5 occupancy)

***Mapping, coverage, and QC***: FastQC (v. 0.11.9) was used to check sequencing quality. Reads were trimmed with TrimGalore (v. 0.6.7) using default parameters and option ‘--paired’. Mapping of the reads from the two replicates was done with BWA^68^ (v. 0.7.17) with default settings to *D. melanogaster* (dm6) genome. Duplicate reads were marked with MarkDuplicates from Picard (v. 3.0.0). The resulting BAM file was sorted and indexed using SAMtools (v. 1.6). Removal of reads with mapping quality below 20, secondary alignment, PCR duplicates, supplementary alignment and reads failing QC was done using SAMtools (v. 1.16.1) to obtain correctly paired end reads with the option ‘samtools view -F 3840 -f 3 -q 20’. Subsequently, reads with a fragment length shorter than 2000 bp were extracted, saved to a BAM file, and indexed using SAMtools (v. 1.16.1). For each CUT&Tag sample, RPKM-normalized signal tracks (bigwig) with 10 bp bins were produced using bamCoverage from deepTools (v. 3.5.1) with options ‘--binSize 10 --exactScaling --extendReads -- ignoreDuplicates --effectiveGenomeSize 125464728 --normalizeUsing RPKM’. To check mapping quality of the reads to their respective genome of the species FastQ Screen (v. 0.15.2) was used with the aligner Bowtie2^69^ (v. 2.4.5) on a subset of 1,000,000 reads. Fragment size for read pairs of each sample was calculated using bamPEFragmentSize from deepTools (v. 3.5.1). Insert size distribution was assessed with Picard CollectInsertSizeMetrics (v. 3.0.0). Checking results from Picard’s MarkDuplicates was done with SAMtools flagstat (v. 1.16.1). Picard CollectAlignmentSummaryMetrics was used to evaluate the mapping quality metrices across samples. The fingerprint (enrichment of signal) was evaluated with deepTools plotFingerprint (v. 3.5.1). Pearson and spearman correlation between samples of RPKM normalized signal tracks were determined using deepTools multiBigwigSummary and plotCorrelation (v. 3.5.1) with a bin size of 1000. QC results were summarised and compared across samples with multiQC (v. 1.13).

The visualisation of SPT5 binding and RNA expression at 2-hour time windows during embryogenesis shown in Fig. 1 was done using pyGenomeTracks^70^ with public RNA-seq data from whole *Drosophila melanogaster* embryos^50^.

***Calculation of the SPT5-based pausing index (SPT5 PI)***: We define the promoter region as TSS ± 250 bp and the gene body region between +250 and the TES. For each gene, we calculated the average SPT5 signal over the entire gene length (promoter plus gene body region, from 250 bp upstream of the TSS to the TES) by summing the RPKM values of the 10-bp bins and dividing by the total number of bins. For each of the five time points analysed, genes with an average SPT5 binding value smaller than 500 RPKM over the entire gene length were considered as not bound by SPT5 and thus excluded from the calculations. The SPT5 signal in the promoter region was determined by the average of the RPKM values from - 250bp to +250bp of the TSS. The signal of the gene body was calculated by summing the RPKM values in the range from +250bp of the TSS to the TES in 10bp bins and dividing by the number of used 10bp bins to obtain the size-normalised SPT5 binding signal of the gene body. The SPT5 pausing index was calculated by dividing the average promoter signal by the size-normalised gene body signal. The calculation of the SPT5 PI was only performed for genes longer than 800bp to ensure an appropriate length of the gene body for the calculation. The categorisation of genes into one of the pause index categories was only performed if the gene was classified as bound to SPT5 during development (average RPKM 500 over the entire gene length) and it was expressed at least in one of the analysed time points (FPKM > 1).

To assess the dynamics of the SPT5 PI across the three clusters shown in Fig. 1, z-scores of the SPT5 PI were calculated for each gene within each cluster (Fig. S2A). Based on the 18–20 h timepoint, genes were divided into two groups: those with a higher SPT5 PI at 18–20 h compared to other timepoints (positive z-score), and those with a lower SPT5 PI at 18– 20 h relative to earlier stages of embryogenesis (negative z-score).

To examine the relationship between gene length and transcriptional pausing (Fig S2F), we analysed gene subsets that consistently exhibited similar SPT5 pausing index (PI) values throughout embryogenesis. Genes with a PI < 1 across all developmental stages were defined as lowly paused, those with a PI between 1 and 3 as moderately paused, and genes with a PI > 3 as highly paused.

#### Spike-in CUT&Tag analysis (depletion experiments)

***Mapping, coverage, and QC:*** FastQC (v. 0.11.9) was used to check sequencing quality. Reads were trimmed with TrimGalore (v. 0.6.7) using default parameters and option ‘-- paired’. Read mapping from the two replicates was done with BWA (v. 0.7.17) with default settings to a joint genome of *D. melanogaster* (dm6) and *D. virilis* (r1.07). Duplicate reads were marked with MarkDuplicates from Picard (v. 3.0.0). The resulting bam file was sorted and indexed using SAMtools (v. 1.16.1). Removal of reads with mapping quality below 20, secondary alignment, PCR duplicates, supplementary alignment and reads failing QC was done with SAMtools (v. 1.16.1) to obtain correctly paired end reads with the option ‘samtools view -F 3840 -f 3 -q 20’. Subsequently, reads with a fragment length shorter than 2000 bp were extracted, saved to a BAM file, and indexed using SAMtools (v. 1.16.1). BAM file reads mapping to *D. melanogaster* or *D. virilis* were separated using SAMtools (v. 1.16.1). For each Depletion CUT&Tag sample, RPKM-normalized signal tracks (bigwig) with 10 bp bins were produced using bamCoverage from deepTools (v. 3.5.4) with options ‘--binSize 10 --exactScaling --extendReads --ignoreDuplicates --effectiveGenomeSize (*D. mel:* 125464728, *D. vir:* 206026697)’. Coverage for replicates was then averaged with bigwigCompare from deepTools (v. 3.5.4) with options “--operation mean --binSize 10”. To check mapping quality of the reads to their respective genome of the species FastQ Screen (v. 0.15.2) was used with the aligner bowtie2 (v. 2.4.5) on a subset of 1,000,000 reads. Checking results from Picard’s MarkDuplicates was done with SAMtools flagstat (version 1.16.1). Insert size distribution was assessed with Picard CollectInsertSizeMetrics (v. 3.0.0). Fragment size for read pairs of each sample was calculated using bamPEFragmentSize from deepTools (v. 3.5.4). Picard CollectAlignmentSummaryMetrics (v. 3.0.0) was used to evaluate the mapping quality metrices across samples. The library complexity was evaluated with Picard EstimateLibraryComplexity (v. 3.0.0). The fingerprint (enrichment of signal) was evaluated with deepTools plotFingerprint (v. 3.5.4). Pearson and spearman correlation between samples and antibodies of RPKM normalized signal tracks were determined using deepTools multiBigwigSummary and plotCorrelation with a bin size of 1000. QC results were summarized and compared across samples with multiQC (v. 1.17).

***Spike-in normalisation***: Spike-in normalisation was employed to enable quantitative comparison between conditions. For each sample, the spike-in normalisation factor was calculated based on the number of reads mapped to the spiked-in exogenous species (*D. virilis*) in the corresponding BAM file. For each contrast (*i.e.*, two ’dark control’ replicates versus two ’blue light treated’ replicates for any given antibody), the spike-in factors from all four samples were normalised by dividing their respective values by the geometric mean of the spike-in factors. This normalisation was performed independently for each contrast to maintain comparability between samples using bamCoverage from deepTools (v. 3.5.4) and a custom python script. Spike-in normalised signal tracks (bigwig) for the *D. melanogaster* genome were generated using bamCoverage (v. 3.5.4) from deepTools with the parameters ‘- -binSize 10 --effectiveGenomeSize 125464728 --extendReads --exactScaling -- ignoreDuplicates --normalizeUsing ’None’’ and option ‘--scaleFactor’ with the value corresponding to the sample normalized spike-in factor. Finally, replicate coverage was averaged with deepTools bigwigCompare (v. 3.5.2) with options ‘--operation mean --binSize 10’.

***Calculation of the average SPT5 depletion:*** To quantify the average percentage reduction in SPT5 binding upon blue light exposure, it is important to consider the different levels of occupancy at the TSS and TES, which reflect SPT5’s different functions at these two regions. If the depletion was complete, the difference between the TSS signal and TES signal would be zero. For each depletion time window, we thus calculated the difference between the TSS and TES SPT5 signal in the dark control and the depleted samples. Their ratio was then used to quantify the percentage reduction of their signal across genes using the formula:

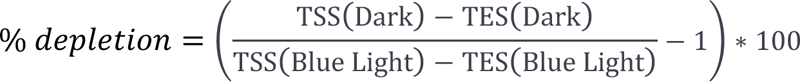

***Peak calling***: For each NELF spike-in CUT&Tag replicate, reads from the BAM file were filtered to retain only paired-end reads with a fragment length < 300 bp using SAMtools (v. 1.16.1). Peak calling was performed with MACS2^71^ (v. 2.2.7.1) using the command callpeak -g 125464728 -p 0.1 -f BAMPE --call-summits. Summits were symmetrically expanded by 200 bp and sorted by signal strength. The expanded summits from both replicates were used as input for IDR^72^ (v. 2.0.4.2) to identify reproducible peaks with the following options: ‘-- input-file-type narrowPeak --rank signal.value --plot -i 0.05 --use-best-multisummit-IDR’. For each reproducible peak, the midpoint was determined and the peak region extended by 500 bp around this centre. Genes were considered bound by NELF if their TSS was located within a reproducible NELF peak.

#### PRO-seq data analysis

***Mapping and coverage:*** Reads were trimmed with Cutadapt (v. 4.1) requiring a minimum base quality of 10 and a minimum adapter length of 20. Trimmed FASTQ files were mapped to the *D. melanogaster* (dm6) reference genome using BWA (v. 0.7.17), and reads with a mapping quality below 30 were removed using SAMtools (v. 1.15.1). Picard’s SortSam (v. 2.18.17) was used to sort the resulting BAM file with the option “SO=coordinate”. Coverage files were generated using the bamCoverage tool from deepTools (v. 3.1.3), using a bin size of 1 bp and normalised to Reads Per Genomic Content (RPGC) to compensate for differences in sequence depth. To remove technical artefacts and unspecific signals, blacklist regions were excluded from the dm6 blacklist^73^.

***PRO-seq pausing index:*** The promoter region was defined as TSS ± 250 bp and the gene body region between +250 and the TES. For each gene, we calculated the average signal over the entire gene length (promoter plus gene body region, from 250 bp upstream of the TSS to the TES) by summing the RPGC values of 1-bp bins and dividing by the total number of bins. The promoter signal was determined by the average RPGC values within the promoter region (TSS ± 250 bp). The gene body signal was calculated by summing RPGC values from +250 bp relative to the TSS to the TES in 1bp bins and dividing by the number of bins to obtain a size-normalised binding signal of the gene body. The PRO-seq pausing index was calculated by dividing the average promoter signal by the size-normalised gene body signal. The PRO-seq pausing index was only calculated for genes longer than 800 bp to obtain a robust metric, similar to the SPT5 PI.

#### Heatmaps and metaprofiles of CUT&Tag and PRO-seq

For the heatmaps after depletion, only expressed genes at the analysed timepoints were considered (FPKM > 1 in a public, time-matched RNA-seq dataset^50^). To refine the analysis, genes shorter than 150 bp as well as tRNA and rRNA genes were excluded. To determine the change of signal upon SPT5 depletion, a log_2_FC of the corresponding signal in the dark and after depletion was obtained using deepTools bigwigCompare (v. 3.5.4) on the merged replicates. To visualize signals from CUT&Tag and PRO-seq datasets at gene loci, deepTools (v3.5.5) was used to generate matrices centered on transcription start sites (TSS), using the parameters --referencePoint TSS, --skipZeros, and --missingDataAsZero. The binSize for the matrices generation was set to 10bp for CUT&Tag and 1bp for PRO-seq datasets.

For the CUT&Tag and PRO-seq metaprofiles and heatmap visualizations, the following procedure was applied to exclude background signals: Bins corresponding to regions beyond the TES of each gene were set to NaN, ensuring they were not included in the calculation of the average signal for the metaprofile. For heatmap visualization, bins beyond the TES were set to zero to prevent the display of signals originating from regions outside the gene.

For the generation of TSS-to-TES scaled metaprofiles, deepTools (v3.5.5) was used to compute matrices with gene bodies scaled to 5 kilobases (kb), while preserving 1 kb of flanking regions upstream of the TSS and downstream of the TES. The command scale-regions was run with the parameters --regionBodyLength 5000, -b 1000, -a 1000, --skipZeros, and --missingDataAsZero and the respective bin size for the experiment to visualize occupancy across gene loci in a standardized manner. To quantify SPT5 depletion across genes, the subtracted signal in the metaprofile plot in Fig 3 were calculated by subtracting the average signal of each 10 bp bin in the blue light condition from the corresponding average signal in the dark condition.

To smooth the signal curves from the PRO-seq visualizations, a Savitzky-Golay filter from SciPy^74^ with a window of 51 data points and a polynomial degree of 3 was used for the TSS-centred metaprofiles, while a window of 501 data points and a polynomial degree of 5 were used for the TSS to TES scaled metaprofiles.

To further assess differences in binding of SPT5, NELF, Pol II pSer2 and Pol II pSer5 at genes which are dysregulated at 18-20 h, following 4h of SPT5 depletion, the binding was assessed in Fig 5 using the 18-20 h, 2 h depletion CUT&Tag and PRO-seq datasets were split into up- and downregulated genes as well as genes which are not dysregulated upon SPT5 depletion.

#### Motif analysis

To investigate promoter motif composition in relation to transcriptional pausing, we selected gene subsets that consistently maintained similar SPT5 promoter index (PI) values throughout embryogenesis. Promoter motifs frequencies were analysed in Fig. S2, using a bootstrap analysis with 10,000 iterations focusing on genes that consistently remained in the same pausing index category. The occurrence of motifs for each gene subset was identified from the CORE database^51^. The CORE database includes a list of core promoter elements/motifs located in the promoter sequences (TSS ±100 bp). In each iteration, 10% of the genes from the three pausing index categories were randomly selected and the frequency of each motif in each category was determined. After bootstrapping, additional Mann-Whitney U tests were performed to identify significant differences in the frequency of motifs between the pausing index categories.

### SUPPLEMENTAL INFORMATION

**Document S1. Figures S1-S4**

**Table S1.** SPT5 binding and RNA expression across five time windows of embryogenesis for genes from the three clusters from Figure1, related to Figure 1

**Table S2.** SPT5 binding across five time windows of embryogenesis for genes from the three SPT5 PI classes, plus list of genes bound by NELF at 3-4, 10-12, and 18-20 h, plus PRO-seq based PI at 3-4 and 18-20 h, related to Figure 2

**Table S3.** Differentially expressed genes from RNA-seq recovery experiments, related to Figure 3

**Table S4.** Differentially expressed genes from RNA-seq after SPT5 depletion in early embryonic stages, related to Figure 4

**Table S5.** Differentially expressed genes from RNA-seq after SPT5 depletion in late embryonic stages, related to Figure 5

## REFERENCES

1. Roeder, R.G. (2019). 50+ Years of Eukaryotic Transcription: an Expanding Universe of Factors and Mechanisms. Nat Struct Mol Biol 26, 783–791. 10.1038/s41594-019-0287-x.

2. Noe Gonzalez, M., Blears, D., and Svejstrup, J.Q. (2021). Causes and consequences of RNA polymerase II stalling during transcript elongation. Preprint at Nature Publishing Group, 10.1038/s41580-020-00308-8 https://doi.org/10.1038/s41580-020-00308-8.

3. Core, L., and Adelman, K. (2019). Promoter-proximal pausing of RNA polymerase II: a nexus of gene regulation. Genes Dev 33, 960. 10.1101/GAD.325142.119.

4. Krebs, A.R., Imanci, D., Hoerner, L., Gaidatzis, D., Burger, L., and Schübeler, D. (2017). Genome-wide Single-Molecule Footprinting Reveals High RNA Polymerase II Turnover at Paused Promoters. Mol Cell 67, 411–422.e4. 10.1016/j.molcel.2017.06.027.

5. Erickson, B., Sheridan, R.M., Cortazar, M., and Bentley, D.L. (2018). Dynamic turnover of paused pol II complexes at human promoters. Genes Dev 32, 1215–1225. 10.1101/gad.316810.118.

6. Steurer, B., Janssens, R.C., Geverts, B., Geijer, M.E., Wienholz, F., Theil, A.F., Chang, J., Dealy, S., Pothof, J., Van Cappellen, W.A., et al. (2018). Live-cell analysis of endogenous GFP-RPB1 uncovers rapid turnover of initiating and promoter-paused RNA Polymerase II. Proc Natl Acad Sci U S A 115, E4368–E4376. 10.1073/PNAS.1717920115.

7. Wada, T., Takagi, T., Yamaguchi, Y., Ferdous, A., Imai, T., Hirose, S., Sugimoto, S., Yano, K., Hartzog, G.A., Winston, F., et al. (1998). DSIF, a novel transcription elongation factor that regulates RNA polymerase II processivity, is composed of human Spt4 and Spt5 homologs. Genes Dev 12, 343–356. 10.1101/gad.12.3.343.

8. Vos, S.M., Farnung, L., Urlaub, H., and Cramer, P. (2018). Structure of paused transcription complex Pol II–DSIF–NELF. Nature 560, 601–606. 10.1038/s41586-018-0442-2.

9. Winston, F., Chaleff, D.T., Valent, B., and Fink, G.R. (1984). Mutations affecting Ty-mediated expression of the HIS4 gene of Saccharomyces cerevisiae. Genetics 107, 179–197. 10.1093/GENETICS/107.2.179.

10. Yamaguchi, Y., Inukai, N., Narita, T., Wada, T., and Handa, H. (2002). Evidence that negative elongation factor represses transcription elongation through binding to a DRB sensitivity-inducing factor/RNA polymerase II complex and RNA. Mol Cell Biol 22, 2918–2927. 10.1128/MCB.22.9.2918-2927.2002.

11. Song, A., and Chen, F.X. (2022). The pleiotropic roles of SPT5 in transcription. Transcription 13, 53–69. 10.1080/21541264.2022.2103366.

12. Bowman, E.A., and Kelly, W.G. (2014). RNA Polymerase II transcription elongation and Pol II CTD Ser2 phosphorylation: A tail of two kinases. Nucleus 5, 224. 10.4161/NUCL.29347.

13. Vos, S.M., Farnung, L., Boehning, M., Wigge, C., Linden, A., Urlaub, H., and Cramer, P. (2018). Structure of activated transcription complex Pol II–DSIF–PAF–SPT6. Nature 2018 560:7720 *560*, 607–612. 10.1038/s41586-018-0440-4.

14. Yamaguchi, Y., Takagi, T., Wada, T., Yano, K., Furuya, A., Sugimoto, S., Hasegawa, J., and Handa, H. (1999). NELF, a Multisubunit Complex Containing RD, Cooperates with DSIF to Repress RNA Polymerase II Elongation. Cell 97, 41–51. 10.1016/S0092-8674(00)80713-8.

15. Yamada, T., Yamaguchi, Y., Inukai, N., Okamoto, S., Mura, T., and Handa, H. (2006). P-TEFb-mediated phosphorylation of hSpt5 C-terminal repeats is critical for processive transcription elongation. Mol Cell 21, 227–237. 10.1016/J.MOLCEL.2005.11.024.

16. Kang, J.Y., Mooney, R.A., Nedialkov, Y., Saba, J., Mishanina, T. V., Artsimovitch, I., Landick, R., and Darst, S.A. (2018). Structural Basis for Transcript Elongation Control by NusG Family Universal Regulators. Cell 173, 1650–1662. 10.1016/J.CELL.2018.05.017.

17. Martinez-Rucobo, F.W., Sainsbury, S., Cheung, A.C.M., and Cramer, P. (2011). Architecture of the RNA polymerase–Spt4/5 complex and basis of universal transcription processivity. EMBO J 30, 1302. 10.1038/EMBOJ.2011.64.

18. Song, A., and Chen, F.X. (2022). The pleiotropic roles of SPT5 in transcription. Transcription 13, 53–69. 10.1080/21541264.2022.2103366.

19. Uzun, Ü., Brown, T., Fischl, H., Angel, A., and Mellor, J. (2021). Spt4 facilitates the movement of RNA polymerase II through the +2 nucleosomal barrier. Cell Rep 36, 109755. 10.1016/j.celrep.2021.109755.

20. Nishimura, K., Fukagawa, T., Takisawa, H., Kakimoto, T., and Kanemaki, M. (2009). An auxin-based degron system for the rapid depletion of proteins in nonplant cells. Nat Methods 6, 917–922. 10.1038/NMETH.1401.

21. Nabet, B., Roberts, J.M., Buckley, D.L., Paulk, J., Dastjerdi, S., Yang, A., Leggett, A.L., Erb, M.A., Lawlor, M.A., Souza, A., et al. (2018). The dTAG system for immediate and target-specific protein degradation. Nature Chemical Biology 2018 14:5 14, 431–441. 10.1038/s41589-018-0021-8.

22. Aoi, Y., Smith, E.R., Shah, A.P., Rendleman, E.J., Marshall, S.A., Woodfin, A.R., Chen, F.X., Shiekhattar, R., and Shilatifard, A. (2020). NELF Regulates a Promoter-Proximal Step Distinct from RNA Pol II Pause-Release. Mol Cell 78, 261–274.e5. 10.1016/J.MOLCEL.2020.02.014.

23. Hu, S., Peng, L., Xu, C., Wang, Z., Song, A., and Chen, F.X. (2021). SPT5 stabilizes RNA polymerase II, orchestrates transcription cycles, and maintains the enhancer landscape. Mol Cell 81, 4425–4439.e6. 10.1016/J.MOLCEL.2021.08.029.

24. Aoi, Y., Takahashi, Y. hei, Shah, A.P., Iwanaszko, M., Rendleman, E.J., Khan, N.H., Cho, B.K., Goo, Y.A., Ganesan, S., Kelleher, N.L., et al. (2021). SPT5 stabilization of promoter-proximal RNA polymerase II. Mol Cell 81, 4413–4424.e5. 10.1016/j.molcel.2021.08.006.

25. Weber, C.M., Ramachandran, S., and Henikoff, S. (2014). Article Nucleosomes Are Context-Specific, H2A.Z-Modulated Barriers to RNA Polymerase. 10.1016/j.molcel.2014.02.014.

26. Chiu, A.C., Suzuki, H.I., Wu, X., Mahat, D.B., Kriz, A.J., and Sharp, P.A. (2018). Transcriptional Pause Sites Delineate Stable Nucleosome-Associated Premature Polyadenylation Suppressed by U1 snRNP. Mol Cell 69, 648. 10.1016/J.MOLCEL.2018.01.006.

27. Fong, N., Sheridan, R.M., Ramachandran, S., and Bentley, D.L. (2022). The pausing zone and control of RNA polymerase II elongation by Spt5: Implications for the pause-release model. Mol Cell 82, 3632–3645.e4. 10.1016/J.MOLCEL.2022.09.001.

28. Martell, D.J., Merens, H.E., Caulier, A., Fiorini, C., Ulirsch, J.C., Ietswaart, R., Choquet, K., Graziadei, G., Brancaleoni, V., Cappellini, M.D., et al. (2023). RNA polymerase II pausing temporally coordinates cell cycle progression and erythroid differentiation. Dev Cell 58, 2112–2127.e4. 10.1016/J.DEVCEL.2023.07.018.

29. Hunt, G., Vaid, R., Pirogov, S., Pfab, A., Ziegenhain, C., Sandberg, R., Reimegård, J., and Mannervik, M. (2024). Tissue-specific RNA Polymerase II promoter-proximal pause release and burst kinetics in a Drosophila embryonic patterning network. Genome Biol 25. 10.1186/S13059-023-03135-0.

30. Saunders, A., Core, L.J., Sutcliffe, C., Lis, J.T., and Ashe, H.L. (2013). Extensive polymerase pausing during Drosophila axis patterning enables high-level and pliable transcription. Genes Dev 27, 1146–1158. 10.1101/GAD.215459.113.

31. Jennings, B.H., Shah, S., Yamaguchi, Y., Seki, M., Phillips, R.G., Handa, H., and Ish-Horowicz, D. (2004). Locus-specific requirements for Spt5 in transcriptional activation and repression in Drosophila. Current Biology 14, 1680–1684. 10.1016/j.cub.2004.08.066.

32. Henriques, T., Scruggs, B.S., Inouye, M.O., Muse, G.W., Williams, L.H., Burkholder, A.B., Lavender, C.A., Fargo, D.C., and Adelman, K. (2018). Widespread transcriptional pausing and elongation control at enhancers. Genes Dev 32, 26–41. 10.1101/gad.309351.117.

33. Aoi, Y., Iravani, L., Mroczek, I.C., Gold, S., Howard, B.C., and Shilatifard, A. (2025). SPT5 regulates RNA polymerase II stability via Cullin 3-ARMC5 recognition. Sci Adv 11, eadt5885. 10.1126/SCIADV.ADT5885/SUPPL_FILE/SCIADV.ADT5885_TABLE_S1.Z IP.

34. Kögler, A.C., Kherdjemil, Y., Bender, K., Rabinowitz, A., Marco-Ferreres, R., and Furlong, E.E.M. (2021). Extremely rapid and reversible optogenetic perturbation of nuclear proteins in living embryos. Dev Cell 56, 2348–2363.e8. 10.1016/j.devcel.2021.07.011.

35. Graveley, B.R., Brooks, A.N., Carlson, J.W., Duff, M.O., Landolin, J.M., Yang, L., Artieri, C.G., Van Baren, M.J., Boley, N., Booth, B.W., et al. (2011). The developmental transcriptome of Drosophila melanogaster. Nature 471, 473–479. 10.1038/nature09715.

36. Zeitlinger, J., Stark, A., Kellis, M., Hong, J.W., Nechaev, S., Adelman, K., Levine, M., and Young, R.A. (2007). RNA polymerase stalling at developmental control genes in the Drosophila melanogaster embryo. Nature Genetics 2007 39:12 *39*, 1512–1516. 10.1038/ng.2007.26.

37. Saunders, A., Core, L.J., Sutcliffe, C., Lis, J.T., and Ashe, H.L. (2013). Extensive polymerase pausing during Drosophila axis patterning enables high-level and pliable transcription. Genes Dev 27, 1146–1158. 10.1101/GAD.215459.113.

38. Lagha, M., Bothma, J.P., Esposito, E., Ng, S., Stefanik, L., Tsui, C., Johnston, J., Chen, K., Gilmour, D.S., Zeitlinger, J., et al. (2013). Paused Pol II coordinates tissue morphogenesis in the Drosophila embryo. Cell 153, 976–987. 10.1016/j.cell.2013.04.045.

39. Pimmett, V.L., Dejean, M., Fernandez, C., Trullo, A., Bertrand, E., Radulescu, O., and Lagha, M. (2021). Quantitative imaging of transcription in living Drosophila embryos reveals the impact of core promoter motifs on promoter state dynamics. Nat Commun 12, 1–16. 10.1038/s41467-021-24461-6.

40. Adelman, K., and Lis, J.T. (2012). Promoter-proximal pausing of RNA polymerase II: emerging roles in metazoans. Nat Rev Genet 13, 720–731. 10.1038/NRG3293.

41. Mahat, D.B., Kwak, H., Booth, G.T., Jonkers, I.H., Danko, C.G., Patel, R.K., Waters, C.T., Munson, K., Core, L.J., and Lis, J.T. (2016). Base-Pair Resolution Genome-Wide Mapping Of Active RNA polymerases using Precision Nuclear Run-On (PRO-seq). Nat Protoc 11, 1455. 10.1038/NPROT.2016.086.

42. Shao, W., and Zeitlinger, J. (2017). Paused RNA polymerase II inhibits new transcriptional initiation. Nat Genet 49, 1045–1051. 10.1038/NG.3867.

43. Hendrix, D.A., Hong, J.W., Zeitlinger, J., Rokhsar, D.S., and Levine, M.S. (2008). Promoter elements associated with RNA Pol II stalling in the Drosophila embryo. Proc Natl Acad Sci U S A 105, 7762–7767. 10.1073/PNAS.0802406105/SUPPL_FILE/0802406105SI.PDF.

44. Bai, S., Fu, K., Yin, H., Cui, Y., Yue, Q., Li, W., Cheng, L., Tan, H., Liu, X., Guo, Y., et al. (2019). The maternal-to-zygotic transition revisited. Development 146. 10.1242/DEV.161471.

45. Harrison, M.M., Marsh, A.J., and Rushlow, C.A. (2023). Setting the stage for development: the maternal-to-zygotic transition in Drosophila. Genetics 225. 10.1093/GENETICS/IYAD142.

46. Blythe, S.A., and Wieschaus, E.F. (2015). Zygotic genome activation triggers the DNA replication checkpoint at the midblastula transition. Cell 160, 1169–1181. 10.1016/j.cell.2015.01.050.

47. Ibarra-Morales, D., Rauer, M., Quarato, P., Rabbani, L., Zenk, F., Schulte-Sasse, M., Cardamone, F., Gomez-Auli, A., Cecere, G., and Iovino, N. (2021). Histone variant H2A.Z regulates zygotic genome activation. Nat Commun 12. 10.1038/S41467-021-27125-7.

48. Weber, C.M., Ramachandran, S., and Henikoff, S. (2014). Article Nucleosomes Are Context-Specific, H2A.Z-Modulated Barriers to RNA Polymerase. 10.1016/j.molcel.2014.02.014.

49. Chiu, A.C., Suzuki, H.I., Wu, X., Mahat, D.B., Kriz, A.J., and Sharp, P.A. (2018). Transcriptional Pause Sites Delineate Stable Nucleosome-Associated Premature Polyadenylation Suppressed by U1 snRNP. Mol Cell 69, 648. 10.1016/J.MOLCEL.2018.01.006.

50. Graveley, B.R., Brooks, A.N., Carlson, J.W., Duff, M.O., Landolin, J.M., Yang, L., Artieri, C.G., Van Baren, M.J., Boley, N., Booth, B.W., et al. (2011). The developmental transcriptome of Drosophila melanogaster. Nature 471, 473–479. 10.1038/nature09715.

51. Dreos, R., Ambrosini, G., Groux, R., Perier, R.C., and Bucher, P. (2017). The eukaryotic promoter database in its 30th year: focus on non-vertebrate organisms. Nucleic Acids Res 45, D51–D55. 10.1093/NAR/GKW1069.

52. Gratz, S.J., Rubinstein, C.D., Harrison, M.M., Wildonger, J., and O’Connor-Giles, K.M. (2015). CRISPR-Cas9 Genome Editing in Drosophila. Curr Protoc Mol Biol 111, 31.2.1-31.2.20. 10.1002/0471142727.MB3102S111.

53. Bonn, S., Zinzen, R.P., Girardot, C., Gustafson, E.H., Perez-Gonzalez, A., Delhomme, N., Ghavi-Helm, Y., Wilczyåski, B., Riddell, A., and Furlong, E.E.M. (2012). Tissue-specific analysis of chromatin state identifies temporal signatures of enhancer activity during embryonic development. Nature Genetics 2012 44:2 44, 148–156. 10.1038/ng.1064.

54. Cavalheiro, G.R., Girardot, C., Viales, R.R., Pollex, T., Ngoc Cao, T.B., Lacour, P., Feng, S., Rabinowitz, A., and Furlong, E.E.M. (2023). CTCF, BEAF-32, and CP190 are not required for the establishment of TADs in early Drosophila embryos but have locus-specific roles. Sci Adv 9. 10.1126/SCIADV.ADE1085/SUPPL_FILE/SCIADV.ADE1085_TABLES_S1_ TO_S5.ZIP.

55. Andrulis, E.D., Guzmán, E., Döring, P., Werner, J., and Lis, J.T. (2000). High-resolution localization of Drosophila Spt5 and Spt6 at heat shock genes in vivo: roles in promoter proximal pausing and transcription elongation. Genes Dev 14, 2635. 10.1101/GAD.844200.

56. Babraham Bioinformatics - FastQC A Quality Control tool for High Throughput Sequence Data https://www.bioinformatics.babraham.ac.uk/projects/fastqc/.

57. Krueger, F., James, F., Ewels, P., Afyounian, E., and Schuster-Boeckler, B. FelixKrueger/TrimGalore: v0.6.7 - DOI via Zenodo. 10.5281/ZENODO.5127899.

58. Dobin, A., Davis, C.A., Schlesinger, F., Drenkow, J., Zaleski, C., Jha, S., Batut, P., Chaisson, M., and Gingeras, T.R. (2013). STAR: ultrafast universal RNA-seq aligner. Bioinformatics 29, 15–21. 10.1093/BIOINFORMATICS/BTS635.

59. Picard Tools - By Broad Institute https://broadinstitute.github.io/picard/.

60. Danecek, P., Bonfield, J.K., Liddle, J., Marshall, J., Ohan, V., Pollard, M.O., Whitwham, A., Keane, T., McCarthy, S.A., and Davies, R.M. (2021). Twelve years of SAMtools and BCFtools. Gigascience 10, 1–4. 10.1093/GIGASCIENCE/GIAB008.

61. Wingett, S.W., and Andrews, S. (2018). FastQ Screen: A tool for multi-genome mapping and quality control. F1000Res 7, 1338. 10.12688/F1000RESEARCH.15931.2.

62. Karolchik, D., Hinrichs, A.S., and Kent, W.J. (2009). The UCSC genome browser. Curr Protoc Bioinformatics Chapter 1. 10.1002/0471250953.BI0104S17.

63. Ramírez, F., Ryan, D.P., Grüning, B., Bhardwaj, V., Kilpert, F., Richter, A.S., Heyne, S., Dündar, F., and Manke, T. (2016). deepTools2: a next generation web server for deep-sequencing data analysis. Nucleic Acids Res 44, W160–W165. 10.1093/NAR/GKW257.

64. Ewels, P., Magnusson, M., Lundin, S., and Käller, M. (2016). MultiQC: summarize analysis results for multiple tools and samples in a single report. Bioinformatics 32, 3047–3048. 10.1093/BIOINFORMATICS/BTW354.

65. Li, B., and Dewey, C.N. (2011). RSEM: Accurate transcript quantification from RNA-Seq data with or without a reference genome. BMC Bioinformatics 12, 1–16. 10.1186/1471-2105-12-323/TABLES/6.

66. Öztürk-Çolak, A., Marygold, S.J., Antonazzo, G., Attrill, H., Goutte-Gattat, D., Jenkins, V.K., Matthews, B.B., Millburn, G., dos Santos, G., Tabone, C.J., et al. (2024). FlyBase: updates to the Drosophila genes and genomes database. Genetics 227. 10.1093/GENETICS/IYAD211.

67. Love, M.I., Huber, W., and Anders, S. (2014). Moderated estimation of fold change and dispersion for RNA-seq data with DESeq2. Genome Biol 15, 1–21. 10.1186/S13059-014-0550-8/FIGURES/9.

68. Li, H. (2013). Aligning sequence reads, clone sequences and assembly contigs with BWA-MEM.

69. Langmead, B., and Salzberg, S.L. (2012). Fast gapped-read alignment with Bowtie 2. Nat Methods 9, 357–359. 10.1038/NMETH.1923.

70. Lopez-Delisle, L., Rabbani, L., Wolff, J., Bhardwaj, V., Backofen, R., Grüning, B., Ramírez, F., and Manke, T. (2021). pyGenomeTracks: reproducible plots for multivariate genomic datasets. Bioinformatics 37, 422–423. 10.1093/BIOINFORMATICS/BTAA692.

71. Zhang, Y., Liu, T., Meyer, C.A., Eeckhoute, J., Johnson, D.S., Bernstein, B.E., Nussbaum, C., Myers, R.M., Brown, M., Li, W., et al. (2008). Model-based analysis of ChIP-Seq (MACS). Genome Biol 9, 1–9. 10.1186/GB-2008-9-9-R137/FIGURES/3.

72. Li, Q., Brown, J.B., Huang, H., and Bickel, P.J. (2011). Measuring reproducibility of high-throughput experiments. 10.1214/11-AOAS466 5, 1752–1779. 10.1214/11-AOAS466.

73. Amemiya, H.M., Kundaje, A., and Boyle, A.P. (2019). The ENCODE Blacklist: Identification of Problematic Regions of the Genome. Sci Rep 9, 1–5.10.1038/S41598-019-45839-Z;TECHMETA=15,23,45;SUBJMETA=114,2785,631;KWRD=COMPUTATIONAL+BIOLOGY+AND+BIOINFORMATICS,GENOME+INFORMATICS.

74. Virtanen, P., Gommers, R., Oliphant, T.E., Haberland, M., Reddy, T., Cournapeau, D., Burovski, E., Peterson, P., Weckesser, W., Bright, J., et al. (2020). SciPy 1.0: fundamental algorithms for scientific computing in Python. Nature Methods 2020 17:3 17, 261–272. 10.1038/s41592-019-0686-2.

